# Morc3 silences endogenous retroviruses by enabling Daxx-mediated H3.3 incorporation

**DOI:** 10.1101/2020.11.12.380204

**Authors:** Sophia Groh, Anna Viktoria Milton, Lisa Marinelli, Cara V. Sickinger, Heike Bollig, Gustavo Pereira de Almeida, Ignasi Forné, Andreas Schmidt, Axel Imhof, Gunnar Schotta

## Abstract

Endogenous retroviruses (ERVs) comprise a significant portion of mammalian genomes. Although, specific ERV loci feature regulatory roles for host gene expression, most ERV integrations are transcriptionally repressed by Setdb1 mediated H3K9me3 and DNA methylation. However, the protein network which regulates deposition of these chromatin modifications is still incompletely understood. Here, we performed a genome-wide sgRNA screen for genes involved in ERV silencing and identified the GHKL ATPase protein Morc3 as top scoring hit. Morc3 knock-out cells display de-repression, reduced H3K9me3 and increased chromatin accessibility of distinct ERV classes. We found that the GHKL ATPase domain of Morc3 is critical for ERV silencing, since mutants which cannot bind ATP, or which are defective in ATP hydrolysis cannot rescue the Morc3 ko phenotype. Proteomic analysis revealed that Morc3 mutant protein which cannot bind ATP fails to interact with the H3.3 chaperone Daxx. This interaction depends on Morc3 SUMOylation, as Daxx lacking the SUMO interaction domain shows reduced association with Morc3. Notably, in Morc3 ko cells, we observed strongly reduced H3.3 on Morc3 binding sites. Thus, our data demonstrate Morc3 as critical regulator of Daxx-mediated H3.3 incorporation to ERV regions.

## INTRODUCTION

Endogenous retroviruses compose a significant portion of mammalian genomes. During evolution, most ERV integrations in mammals were highly mutated or partially deleted and are thus unable to generate functional retroviral particles. However, ERV LTRs harbor binding sites for transcription factors and can act as regulatory elements to drive host gene expression (Thompson et al., 2016). This physiological role for regulation of normal development is contrasted by pathological effects of aberrant ERV regulation in diseases (Deniz et al., 2020).

A prominent silencing pathway for many ERV families involves establishment of H3K9me3 and DNA methylation (Groh and Schotta, 2017). How these modifications mediate silencing is less understood. Probably, the low accessibility associated with this chromatin state could prevent transcription factor binding or RNA Polymerase activity. The current model for establishment of H3K9me3 on ERVs is based on sequence-specific binding of KRAB-ZnF proteins, which recruit the co-repressor Trim28 and the histone methyltransferase Setdb1 (Geis and Goff, 2020). The establishment of DNA methylation is likely to happen during early embryogenesis, and maintenance of high methylation levels on ERVs is later ensured by Uhrf1/Dnmt1 (Ramesh et al., 2016; Walsh et al., 1998). Full establishment of H3K9me3 and DNA methylation as well as reduced chromatin accessibility on ERVs require activities, such as H3.3 deposition by Atrx/Daxx (Elsasser et al., 2015; He et al., 2015; Sadic et al., 2015; Wasylishen et al., 2020), chromatin remodeling by Smarcad1 (Sachs et al., 2019) and chromatin assembly by Chaf1a/b (Yang et al., 2015). Currently, it is not clear how these activities are coordinated to restrict chromatin accessibility and to mediate heterochromatin spreading on ERVs.

Genome-wide siRNA/sgRNA screens based on retroviruses like MMLV and MSCV have revealed genes that also play critical roles in ERV silencing (Fukuda et al., 2018; Yang et al., 2015). However, no equivalent screens were performed using endogenous retroviral sequences. We have previously identified a small heterochromatin inducing sequence (SHIN) in Intracisternal A Type particle (IAP) elements which triggers strong reporter silencing (Sadic et al., 2015). Using a genome-wide sgRNA screen for genes involved in SHIN silencing we identified Morc3 as novel player in ERV silencing. Morc3 is a GHKL type ATPase protein which can form a closed dimer in the ATP bound state (Li et al., 2016). It further contains a CW type zinc finger domain that negatively regulates ATPase activity (Andrews et al., 2016). Ligands of the CW domain, e.g. histone H3K4me3 or influenza virus protein NS1 peptides relieve suppression of ATPase activity and could regulate the turnover of ATPase domain mediated dimerization and opening (Zhang et al., 2019a; Zhang et al., 2019b). Although H3K4me3-mediated interaction of Morc3 with promoter regions in mouse ES cells was reported (Li et al., 2016), the molecular roles of Morc3 in transcriptional regulation or chromatin organization are currently unclear. Our data reveal Morc3 as critical player in ERV silencing. We demonstrate that Morc3 binds ERV sequences and, that loss of Morc3 results in increased chromatin accessibility, reduced H3K9me3 and de-repression of ERVs. We detect an interaction of Morc3 with the H3.3 chaperone Daxx, that depends on the Morc3 ATPase cycle and SUMO-interaction by Daxx. This interaction is crucial for Daxx-mediated H3.3 incorporation as Morc3 ko ES cells lose H3.3 on ERVs. Thus, our data indicate Morc3 as novel regulator of Daxx-mediated H3.3 incorporation.

## RESULTS

### Identification of Morc3 as a novel ERV silencing factor

To identify novel factors regulating IAPEz silencing we performed a genome-wide sgRNA screen based on the integration of an IAP SHIN reporter in mouse ES cells (T90 cells, Figure 1A). This reporter contains a doxycycline inducible promoter driving the expression of EGFP and a zeocin resistance gene. In the wild type condition, doxycycline induction results in poor reporter activation due to its heterochromatic state, whereas impaired SHIN silencing allows doxycycline induced reporter activity (Sadic et al., 2015). We used a pooled genome wide lentiviral sgRNA library (Sanjana et al., 2014) to transduce T90 ES cells, followed by doxycycline induction to activate the reporter locus. Reporter activity also results in zeocin resistance. Therefore, we applied zeocin selection to detect cells with an activated reporter resulting from impaired heterochromatin. As control we collected a second pool of cells without selection pressure. We then identified the sgRNAs which were enriched in the zeocin selected cells (Figure 1B, Supplementary Table 1). Notably, top enriched sgRNAs targeted the major known ERV silencing factors, such as Dnmt1, Uhrf1, Setdb1, Trim28 and Atrx/Daxx (Figure 1B). Another top hit was Morc3, a protein not previously implicated in ERV silencing. Due to the mode of selection with zeocin, we also identified genes which might be related to zeocin resistance, DNA damage repair and apoptosis (Supplementary Table 1). To validate selected screening hits, we performed SHIN silencing maintenance assays for selected candidate gene knockouts (Figure 1C). Control sgRNA treatment did not result in an activatable SHIN reporter, whereas sgRNA knock-out of Setdb1 and Morc3 resulted in significant SHIN de-repression (Figure 1D). We further tested additional screening candidates in the SHIN maintenance assay and could partially confirm our screening results (Figure 1E). We then complemented the functional testing with a SHIN initiation silencing assay in which we test whether a newly introduced SHIN reporter with constitutively active promoter can be silenced in different genetic backgrounds (Figure 1F). Here, the results differed from the maintenance assay as the DNA methylation pathway appeared less important for establishing silencing, while Setdb1/Trim28 and Atrx/Daxx were still critical for silencing (Figure 1G). Since Morc3 was important in both maintenance and initiation of silencing we decided to functionally characterize this protein in more detail.

**Figure 1.**
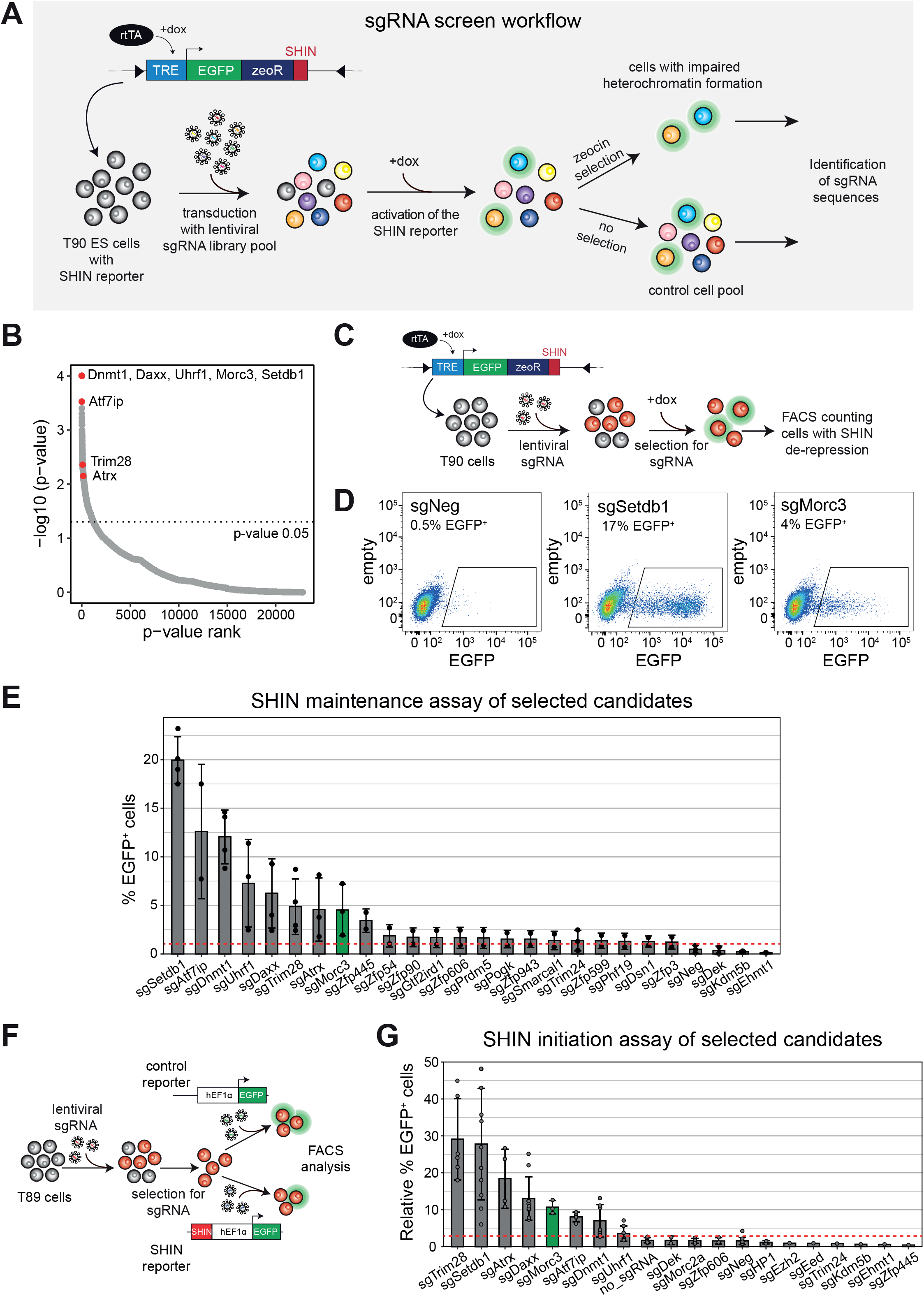
A genome-wide sgRNA screen for SHIN silencing identifies Morc3. (A) Setup of the sgRNA screen. T90 ES cells containing an inducible SHIN reporter (Sadic et al., 2015) were transduced with a genome-wide sgRNA. Cells were selected for sgRNA vectors and the reporter was activated with doxycycline. Two independent cell batches were cultured to harvest non-selected cells representing the control and zeocin-resistant cells representing cells with impaired heterochromatin on the SHIN reporter. (B) Dot plot showing the sgRNA screen results ordered by p value rank. Major ERV silencing factors are indicated. Morc3 represents a top hit in the screen. (C) Schematic of the SHIN silencing maintenance assay. T90 ES cells were transduced with sgRNAs for selected candidates and treated for integration with puromycin. Subsequently, doxycycline was added to the cells to induce reporter activity. EGFP expression was analyzed by FACS. (D) FACS plots depicting EGFP fluorescence in cells with activated SHIN reporter. Almost no activity was detected in cells transduced with a control sgRNA, demonstrating full SHIN silencing. sgRNAs targeting Setdb1 or Morc3 result in SHIN de-repression as indicated by cells showing EGFP expression. (E) Bar plot depicting the results of the SHIN maintenance silencing assay with selected candidate genes. The red dotted line indicates the background reporter activity. (F) Schematic of the SHIN silencing initiation assay. T89 ES cells were transduced with sgRNAs for selected candidates and treated for integration with puromycin. Subsequently, the cells were transduced with a control virus without the SHIN sequence and the SHIN reporter with constitutive promoter, respectively. EGFP expression was analyzed by FACS and the percentage of EGFP expressing cells, relative to the control reporter was calculated. (G) Bar plot depicting the results of the SHIN initiation silencing assay with selected candidate genes. The red dotted line indicates the background reporter activity.

### Morc3 binds different families of endogenous retroviruses

To assess if Morc3 directly regulates ERV silencing we performed ChIP-seq experiments using a knock-in cell line that expresses 3xFLAG tagged Morc3 at endogenous levels (Supplementary Figure 1). We identified 2737 peaks which were shared between two replicates (Supplementary Table 2). Annotation with genomic features revealed that most Morc3 peaks associate with ERVs (Figure 2A). We then classified the ERV families to which Morc3 peaks associate and found a major enrichment with IAPEz, LTRIS2 and RLTR families (Figure 2B). Due to lack of polymorphisms many ERV integrations cannot be precisely mapped. To better assess association of Morc3 with ERV families we used RepEnrich2 (Criscione et al., 2014) to categorize all mappable Morc3 ChIP-seq reads into ERV families. This analysis confirmed prominent association of Morc3 with IAP, RLTR and LTRIS families (Figure 2C, Supplementary Table 3). The most prominent ERV class bound by Morc3 are IAPEz elements, from which the SHIN sequence used in our screen originates. We generated cumulative coverage plots for IAPEz elements to determine whether Morc3 displays preferential binding with distinct IAPEz features. Interestingly, comparison of the Morc3 profile with ChIP-seq profiles for Setdb1, Trim28 and H3K9me3 (Figure 2D) revealed prominent co-enrichment of the silencing factors with the IAP GAG region containing the SHIN sequence and with the 5’UTR region which is also able to induce reporter silencing (Rowe et al., 2010). The similarity of the Morc3 binding pattern to Setdb1 and Trim28 suggests a functional interplay with the H3K9me3 pathway (Figure 2D). To investigate whether Morc3 more generally associates with Setdb1 and Trim28 on its identified peaks, we generated a read-density heatmap of Setdb1, Trim28 and H3K9me3 on Morc3 peaks (Figure 2E). This analysis also revealed prominent enrichment of Setdb1 and Trim28 on Morc3 peaks, further supporting Morc3 association with H3K9me3 repressed heterochromatin.

**Figure 2.**
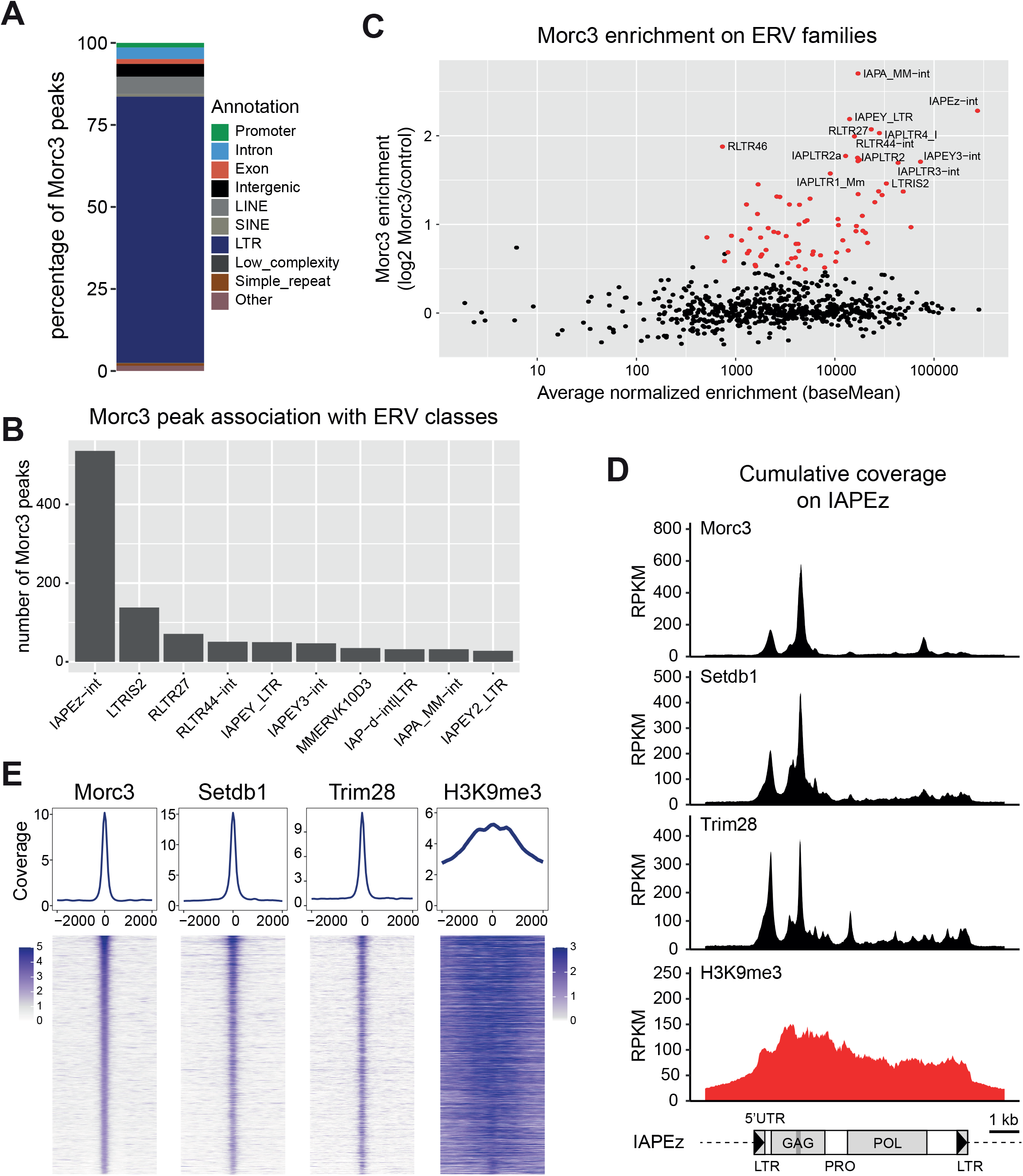
Morc3 binds to different families of endogenous retroviruses. (A) Annotation statistics of Morc3 peaks. Most peaks are on ERVs. (B) Morc3 peak association with ERV families. Bar plot shows the number of Morc3 peaks which associate with the ten most enriched ERV families. (C) Dot plot showing Morc3 ChIP enrichment on ERV families. Red dots highlight ERV families with significant Morc3 over background enrichment. Selected ERV families are labeled. (D) Cumulative coverage plot of Morc3, Setdb1, Trim28 and H3K9me3 ChIP-seq profiles on IAPEz elements. Prominent enrichment is over the 5’UTR and the GAG region. The position of the SHIN sequence is indicated as dark gray bar. (E) Morc3 peaks are associated with Setdb1 and Trim28 binding and feature high H3K9me3. Top panel: Density plot showing the average occupancy of Morc3, Setdb1, Trim28 and H3K9me3 on Morc3 peaks. Lower panel: Read-density heat map showing the normalized coverage of Morc3, Setdb1, Trim28 and H3K9me3 on Morc3 binding sites. Distance from the peak center is given in bp.

### Morc3 knock-out results in ERV de-repression

To investigate the role of Morc3 in ERV silencing we generated Morc3 knock-out ES cells (Supplementary Figure 2). Transcriptome analysis by RNA-seq revealed significant dys-regulation of 252 genes (adjusted p-value < 0.01) with a trend towards genes being up-regulated (Figure 3A, Supplementary Table 4). We then asked if Morc3 could be directly involved in regulation of these genes by determining Morc3 peaks in the vicinity of their transcription start sites. We found that 64 up-regulated and 18 down-regulated genes had Morc3 peaks in < 100kb distance to their promoter (Supplementary Table 5). When we investigated the detailed peak annotation of these Morc3 peaks we found strong association with ERV LTR sequences. The highest enriched ERV class was LRTIS2, which was found on 25 Morc3 peaks which associate with up-regulated genes (Supplementary Table 5). These data suggest that genes are indirectly regulated through de-repressed ERV LTRs which could act as enhancer elements for neighboring genes. Three prominent examples for up-regulated genes are shown in Figure 3B. Irak3 has two Morc3 peaks in its gene body of which one associates with ORR1A3/Lx9 repeats. Ube2l6 features Morc3 binding in an LTRIS2 element, overlapping the 3’UTR region. Cd200 shows Morc3 binding with an LTRIS2 element down-stream of its gene body. To attribute the effects on gene regulation to Morc3 function, we generated rescue ES cell lines by transducing the full length Morc3 cDNA into Morc3 ko ES cells (Supplementary Figure 2A). We then quantified expression of selected Morc3 target genes by RT-qPCR in wild type, Morc3 ko and rescue cells. All selected target genes displayed de-repression in Morc3 ko, whereas expression was reduced in rescue cells (Figure 3C).

**Figure 3.**
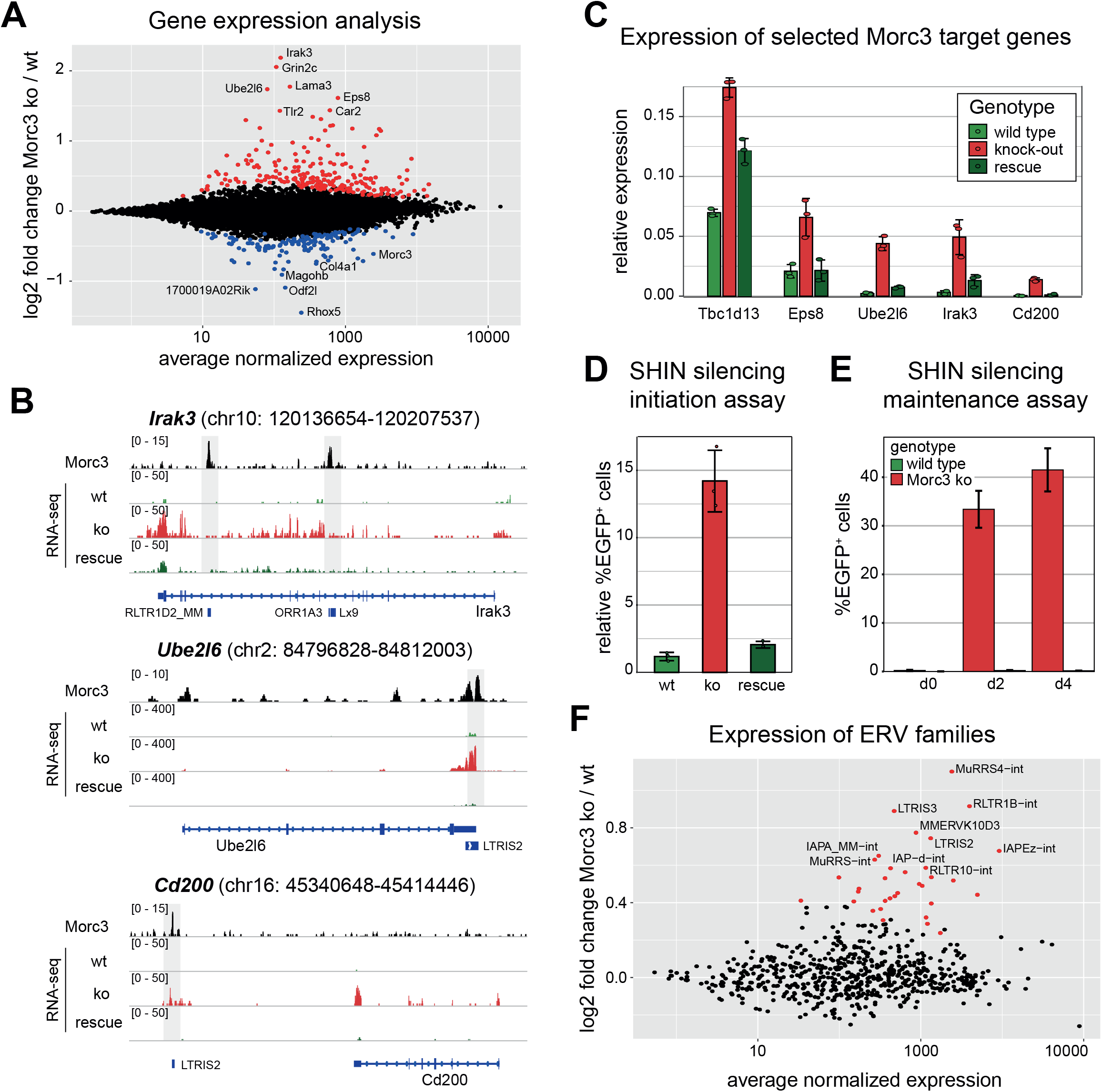
Morc3 knock-out cells display de-repression of genes and distinct ERV families. (A) Dot plot showing average expression vs. log2-fold change of coding genes in wild type vs. Morc3 knock-out ES cells. Colored dots indicate genes with significantly changed expression (adjusted p-value < 0.05, n=3 for each condition). Positions of relevant genes are indicated. (B) Genome browser view of Morc3-dependent expression changes on selected target genes (Irak3, Ube2l6 and Cd200). Morc3 peaks are located on ERVs in close vicinity to these genes. Transcriptional up-regulation of these target genes in Morc3 ko cells can be rescued by expression of wild type Morc3. (C) RT-qPCR analysis of selected Morc3 target genes in wild type, Morc3 ko and Morc3 rescue ES cells. Bar graph depicts relative expression to control genes (Actin and Hprt). Error bars indicate standard deviation of replicate experiments (n=3). Individual data points are shown as colored dots. (D) SHIN initiation silencing assay in wild type, Morc3 ko and Morc3 rescue ES cells. Bar graph depicts relative percentage of EGFP positive cells of SHIN-reporter transduced cells relative to control virus transduced cells. Error bars indicate standard deviation of replicate experiments (n=3). Individual data points are shown as colored dots. (E) SHIN maintenance silencing assay in wild type and Morc3 ko ES cells. Bar graph depicts percentage of EGFP positive cells after doxycycline induction for two and four days, respectively. Error bars indicate standard deviation of replicate experiments (n=3). (F) Dot plot showing average expression vs. log2-fold change of ERV families in wild type vs. Morc3 knock-out ES cells. Colored dots indicate genes with significantly changed expression (adjusted p-value < 0.05, n=3 for each condition). Positions of relevant ERV families are indicated.

To determine Morc3 roles in ERV repression we performed SHIN silencing initiation assays. Morc3 knock-out cells displayed impaired SHIN silencing, whereas rescue cells can efficiently establish SHIN repression (Figure 3D). We also generated Morc3 knock-out cells from T90 SHIN reporter ES cells to investigate Morc3-dependent silencing maintenance (Supplementary Figure 2B). Without doxycycline induction, no EGFP expression could be detected in wild type or Morc3 ko cells (Figure 3E). We then monitored EGFP expression from the SHIN reporter with two and four days of doxycycline induction. Wild type cells failed to activate the reporter and almost no EGFP positive cells could be detected (Figure 3E). In contrast, silencing of the SHIN reporter was strongly impaired in Morc3 ko cells (Figure 3E), demonstrating that Morc3 is important for the maintenance of SHIN silencing.

To determine if IAP and potentially other ERV families are de-repressed in Morc3 ko cells, we counted RNA-seq reads corresponding to ERVs by RepEnrich2 and calculated differentially expressed ERV families (Figure 3F, Supplementary Table 6). We found that indeed, major Morc3 target families, such as IAP, RLTR and LTRIS displayed significant de-repression, although the extent of regulation is greatly reduced compared to Setdb1 knock-out where IAP de-repression is around 10-100-fold (Karimi et al., 2011; Matsui et al., 2010). These data suggest that Morc3 is a contributing factor for ERV silencing which acts redundantly with other mechanisms in the context of ERVs. However, individual ERV integrations, probably with less redundancy in silencing mechanisms, are strongly de-repressed in Morc3 ko ES cells and affect the expression of neighboring genes.

### Morc3 ko results in reduced H3K9me3 and increased chromatin accessibility

To characterize the chromatin changes coinciding with Morc3 loss, we measured H3K9me3 in wild type, Morc3 ko and rescue cells. Visual inspection of H3K9me3 profiles on Morc3 peaks revealed different patterns of changes. Therefore, we clustered Morc3 peaks according to changes in H3K9me3 patterns and generated a read-density heatmap of Morc3 peak clusters with H3K9me3 pattern in wild type cells, together with fold change representation in Morc3 ko and rescue cells (Figure 4A). We found that H3K9me3 was generally reduced in cluster I. Clusters II and III displayed a biased reduction of H3K9me3 towards one side of the peak center. Clusters IV and V showed reduced H3K9me3 mainly in the peak center and cluster VI did not show appreciable changes in H3K9me3. The genome browser screenshot for the Morc3 target gene *Irak3* displays two Morc3 peaks representing different clusters (Figure 4C). One peak which is most distal to the Irak3 promoter features H3K9me3, which is not changed in Morc3 ko cells. The promoter proximal peak displays a one-sided reduction of H3K9me3 from the peak center extending towards downstream (Figure 4C).

**Figure 4.**
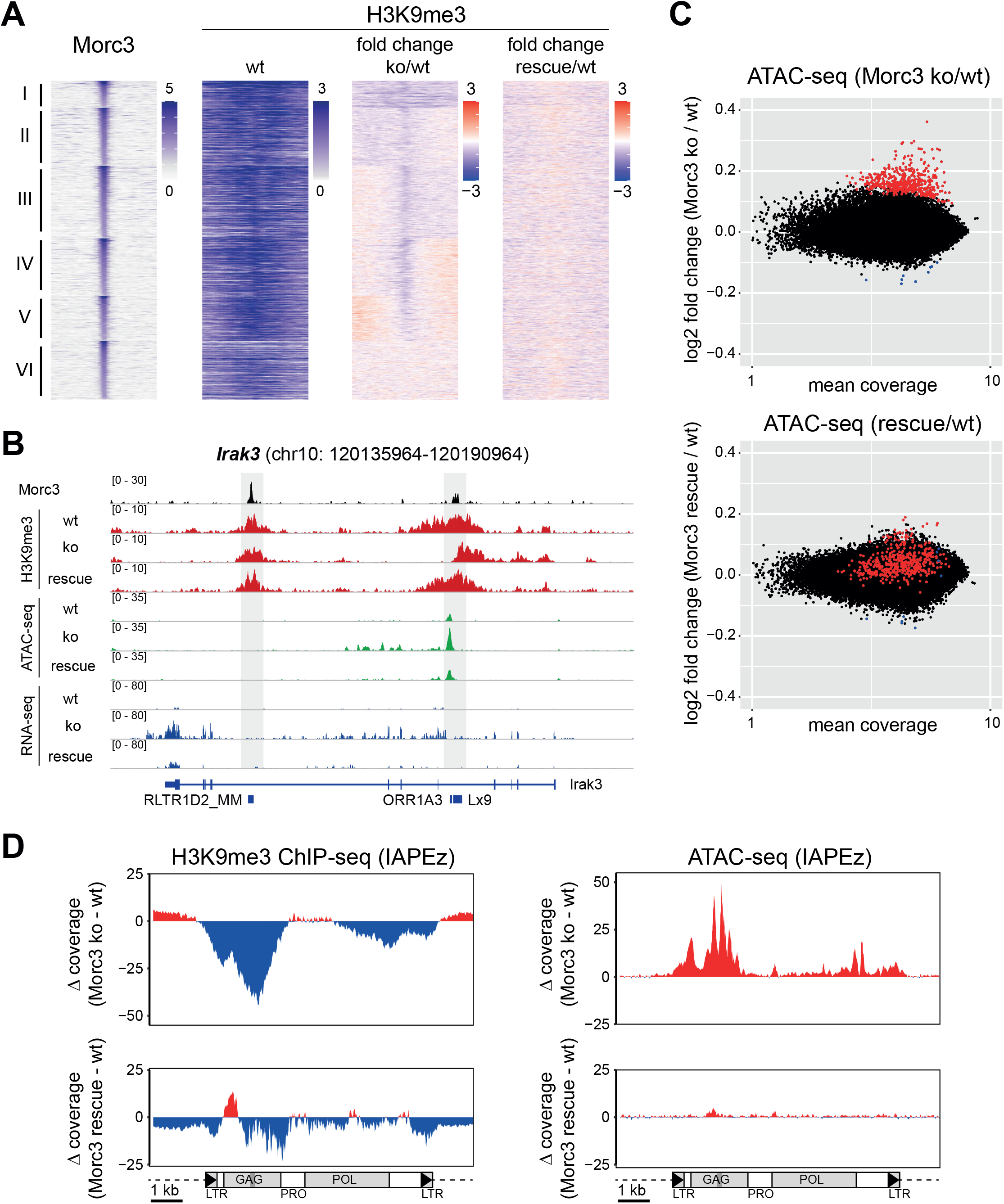
Morc3 knock-out results in reduced H3K9me3 and increased chromatin accessibility on target regions. (A) Changes in H3K9me3 on Morc3 peaks. Morc3 peaks were grouped in six clusters according to changes in H3K9me3 in Morc3 ko ES cells. Read density heatmaps show the normalized coverage of Morc3 and H3K9me3 on Morc3 peaks (left columns), the fold change in H3K9me3 signal between Morc3 ko and wild type ES cells (third column) and the fold change in H3K9me3 between Morc3 rescue and wild type cells (fourth column). Depletion of H3K9me3 is colored in blue, increased H3K9me3 appears red. Classes I-V display specific reduction in H3K9me3 in a larger region (class I-III) or more selective in the peak center (class IV-V). H3K9me3 is restored close to wild type levels in Morc3 rescue cells. (B) Genome browser view of Morc3-dependent chromatin changes on an example target gene (Irak3). Positions of two Morc3 peaks are indicated by gray boxes. The right Morc3 binding site displays selective loss of H3K9me3 and increased chromatin accessibility, concomitant with elevated transcription. Chromatin and transcriptional changes are rescued in Morc3 rescue ES cells. (C) Dot plot showing average coverage vs. log2-fold change of ATAC peaks in wild type vs. Morc3 knock-out ES cells (top panel) and wild type vs. Morc3 rescue ES cells (lower panel). Colored dots indicate ATAC peaks with significantly increased (red dots) or decreased (blue dots) coverage in Morc3 ko ES cells (adjusted p-value < 0.05, n=3 for each condition). ATAC coverage of these peaks is largely normalized in Morc3 rescue ES cells. (D) Reduced H3K9me3 and increased chromatin accessibility on IAPEz elements. Plots display the difference in cumulative coverage between Morc3 ko and wild type ES cells (top panel) and Morc3 rescue and wild type ES cells (lower panel) for H3K9me3 and ATAC-seq. Reduced H3K9me3 and increased accessibility is mainly in the 5’UTR and GAG region. The position of the SHIN sequence is indicated as gray bar.

H3K9me3 heterochromatin is characterized by low chromatin accessibility which could prevent efficient access of transcriptional activators (Becker et al., 2016; Soufi et al., 2012). To test if chromatin accessibility changes in Morc3 ko cells, we performed ATAC-seq experiments in wild type, Morc3 ko and rescue ES cells. Altogether we identified 444 peaks with significantly higher accessibility in Morc3 ko ES cells, while only 10 peaks were less accessible (Figure 4B). Increased chromatin accessibility is directly related to Morc3 loss as we found reduced chromatin accessibility of these peaks in Morc3 rescue cells (Figure 4B). Interestingly, up-regulated ATAC peaks were found in the vicinity of Morc3 peaks, but usually not directly overlapping with the Morc3 peak center. An example for this situation is shown in the genome browser screenshot for Irak3 (Figure 4C), where the promoter proximal Morc3 peak displays increased ATAC-seq coverage across a larger genomic interval flanking the peak center. Other Morc3 targets display similar chromatin changes (Supplementary Figure 3), indicating that Morc3 is required to repress transcriptional activators from binding to ERV sequences. Loss of Morc3 leads to reduced H3K9me3 and increased chromatin accessibility, most likely coinciding with transcription factor binding, which, in turn, results in gene upregulation.

Since IAPEz elements represent the major target of Morc3 peaks, we calculated the difference in H3K9me3 and ATAC-seq signals comparing Morc3 ko vs. wild type and rescue vs. wild type cells (Figure 4D). We found that H3K9me3 is decreased in the IAP-GAG region, concomitant with increased chromatin accessibility (Figure 4D, top panels). These chromatin changes were reverted in Morc3 rescue cells (Figure 4D, lower panel). We extended this analysis using RepEnrich2 analysis and detected reduced H3K9me3 and increased chromatin accessibility on other ERV families (Supplementary Figure 4). Together, our data demonstrate that Morc3 is critical for the full maintenance of H3K9me3 and low chromatin accessibility on distinct ERV families.

### An efficient ATPase cycle is critical for Morc3-dependent ERV silencing

Morc3 contains a C-terminal coiled coil domain which mediates dimerization; however, the N-terminal GHKL ATPase domain can only dimerize in an ATP bound state (Figure 5A). ATP hydrolysis and dissolution of the ATPase domain dimer is negatively regulated by the CW domain. Ligand binding of the CW domain relieves this inhibition and results in higher ATPase activity (Zhang et al., 2019b). To test the function of the Morc3 ATPase cycle for ERV silencing we generated distinct Morc3 mutant rescue cell lines (Figure 5A, Supplementary Figure 5A). The ATP hydrolysis mutant (E35A) is impaired to dissolve the ATPase dimer, the ATP binding mutant (G101A) is unable to dimerize the ATPase domains and, the CW ligand binding mutant (W419A) has a reduced ATPase activity, resulting in a slowed-down ATPase cycle. All rescue cell lines display a similar nuclear distribution of Morc3 (Figure 5B), which is consistent with previous reports (Mimura et al., 2010). ChIP-seq analyses for the Morc3 mutant proteins revealed concordant localization with Morc3 peaks and IAPEz elements (Supplementary Figure 5B,C), with the exception of the ATP hydrolysis mutant, which failed to properly localize to Morc3 targets (Supplementary Figure 5B,C).

**Figure 5.**
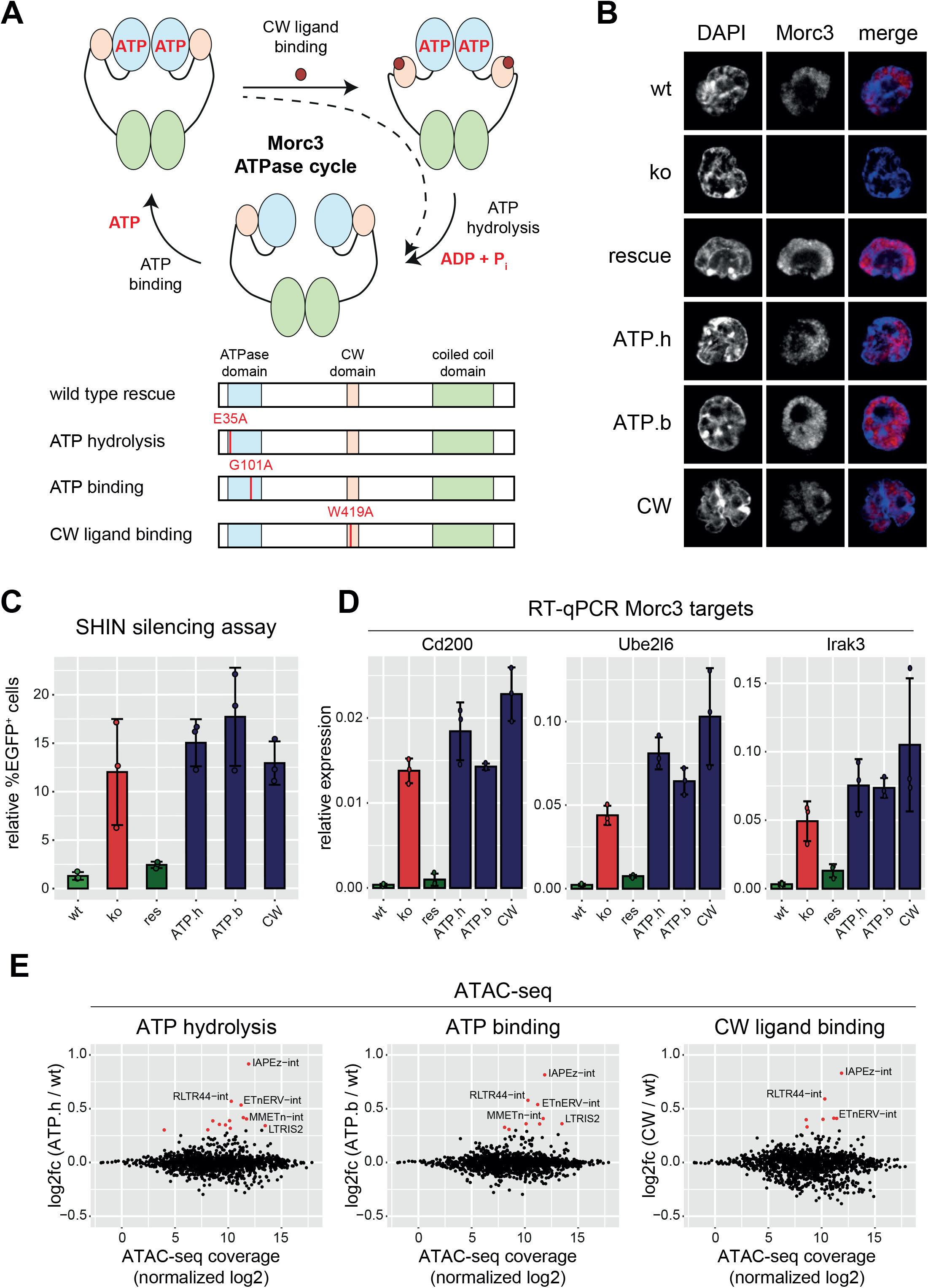
Morc3 silencing activity depends on a functional ATPase cycle. (A) Morc3 mutations which impair the ATPase cycle. Three mutant constructs were generated to impair ATP binding, ATP hydrolysis and CW ligand interaction (top panel). These mutant proteins affect different aspects of the Morc3 ATPase cycle: ATP binding is needed for dimerization of the ATPase domain. The CW domain is a negative regulator of the ATPase activity and requires ligand interaction to dissociate from the ATPase domain. Subsequent ATP hydrolysis results in dissolution of the ATPase domain dimer. The ATP binding mutant (ATP.b) is not able to dimerize; the ATP hydrolysis mutant (ATP.h) is constitutively dimeric and the CW mutant (CW) results in reduced ATPase activity resulting in a slower ATPase cycle. (B) Immunofluorescence analysis of wild type and Morc3 mutant rescue cells. In wild type cells, Morc3 displays a broad nuclear staining, which is lost in Morc3 knock-out cells. All Morc3 mutant proteins display a similar staining pattern as the wild type protein. (C) SHIN initiation silencing assay with Morc3 mutant rescue cell lines. Bar graph depicts relative percentage of EGFP positive cells of SHIN-reporter transduced cells relative to control virus transduced cells. Error bars indicate standard deviation of replicate experiments (n=3). Individual data points are shown as colored dots. Data for wild type, ko and rescue are from figure 3D for comparison. (D) RT-qPCR analysis of selected Morc3 target genes in Morc3 mutant rescue ES cells. Bar graphs depict relative expression to control genes (Actin and Hprt). Error bars indicate standard deviation of replicate experiments (n=3). Individual data points are shown as colored dots. Data for wild type, ko and rescue are from figure 3C for comparison. (E) Dot plot showing average ATAC-seq coverage vs. log2-fold change of ERV families in wild type vs. Morc3 mutant rescue ES cells (ATP hydrolysis mutant, ATP binding mutant, CW ligand binding mutant). Colored dots indicate ATAC peaks with significantly increased (red dots) or decreased (blue dots) coverage in Morc3 mutant rescue ES cells (adjusted p-value < 0.05, n=3 for each condition).

Next, we functionally tested the Morc3 mutant rescue cells in SHIN initiation silencing assays and found that all mutant proteins failed to induce SHIN silencing (Figure 5C). We also performed RT-qPCR analysis for selected Morc3 target genes and observed de-repression, comparable to Morc3 knock-out cells (Figure 5D). Finally, we performed ATAC-seq experiments with the mutant rescue cell lines to estimate Morc3-dependent chromatin changes. All three mutants displayed increased accessibility of ERV families, only the CW mutant appeared slightly less compromised (Figure 5E). Difference in cumulative ATAC-seq coverage on IAPEz elements demonstrated compromised chromatin architecture of this ERV family in Morc3 mutant rescue cells (Supplementary Figure 5D). Consistent with the de-repression of Morc3 target genes we detected increased chromatin accessibility on associated Morc3 peaks (Supplementary Figure 6). Together our data demonstrate that a fully functional ATPase cycle is critical for Morc3 functionality.

### Morc3 regulates Daxx-mediated H3.3 incorporation

Conformational changes during the Morc3 ATPase cycle might affect interactions with other proteins, that could be related to ERV silencing. To test this hypothesis, we determined the protein interaction context of Morc3 using ChIP mass-spectrometry analysis. This approach uses crosslinking and solubilization of chromatin through sonication. Therefore, proteins detected in this approach represent direct and stable protein interactions as well as low affinity interactions which are more transient in nature and proteins for which the interaction with Morc3 is mediated through DNA/RNA fragments. We detected 489 proteins which significantly associate with Morc3 (Figure 6A, Supplementary Table 7). Importantly, we detect the major ERV silencing factors Setdb1, Trim28, Atrx/Daxx and Dnmt1/Uhrf1. We also observe strong association with SUMO proteins, corresponding to high SUMOylation of Morc3 (Mimura et al., 2010) and consistent with the role of SUMO in ERV repression (Yang et al., 2015). The functional categorization of Morc3 interactors using Panther protein class enrichment analysis revealed that many proteins in the Morc3 context belong to chromatin and histone-modifying activities (Figure 6B). Interestingly a large proportion of proteins related to RNA binding and RNA processing, suggesting that Morc3 may have roles in RNA metabolism.

**Figure 6.**
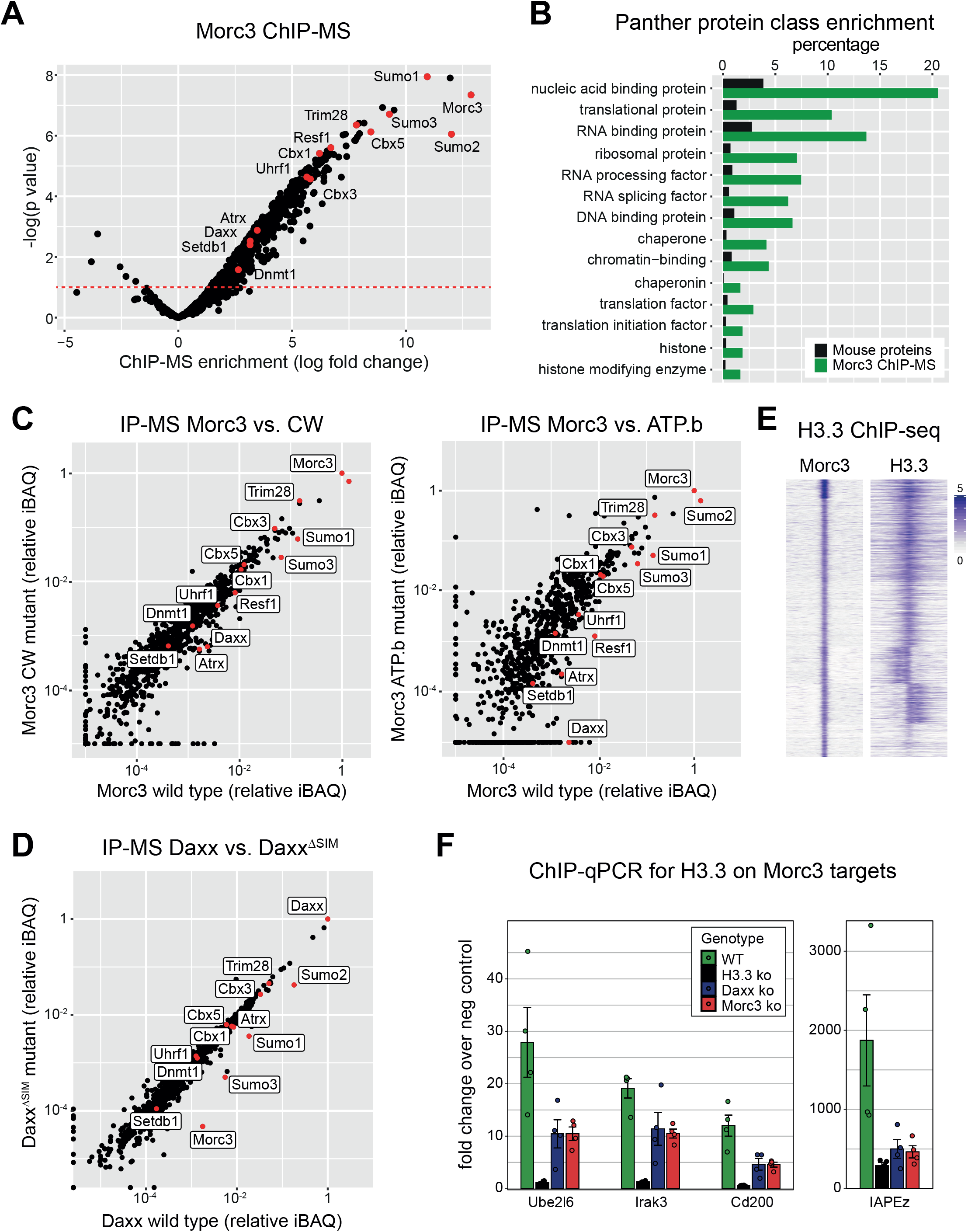
Morc3 dimerization is needed for Daxx recruitment. (A) ChIP-MS identification of Morc3-associated proteins. Morc3 ChIP-MS experiments (n=3) were performed with Morc3-3xFLAG knock-in ES cells and wild type ES cells using FLAG antibody. Dot plot shows log fold change enrichment vs. −log(p value) of proteins in Morc3-3xFLAG vs wild type (background IP) cells. Positions of labelled proteins are indicated by red dots. (B) Panther protein class enrichment analysis of proteins which are significantly enriched in the Morc3 ChIP-MS. Bar graph shows the percentage of total mouse proteins (black) or Morc3 ChIP-MS proteins (green) in significantly enriched Panther protein classes (p value < 0.05). (C) ChIP-MS analysis of wild type vs. mutant Morc3 proteins. Dot plot shows relative iBAQ values of proteins quantified in wild type vs. Morc3 CW mutant (CW, left panel) and wild type vs. Morc3 ATP binding mutant (ATP.b, right panel). Positions of ERV regulators are indicated. (D) ChIP-MS analysis of Daxx wild type vs. Daxx^ΔSIM^ mutant proteins. Dot plot shows relative iBAQ values of proteins quantified in Daxx rescue cell lines expressing wild type Daxx or Daxx^ΔSIM^ mutant protein. Positions of ERV regulators are indicated by red dots. (E) H3.3 is enriched on Morc3 peaks. Read-density heat map showing the normalized coverage of Morc3 and H3.3 on Morc3 peaks. (F) Morc3 is critical for H3.3 deposition. ChIP-qPCR for H3.3 on selected Morc3 target regions in wild type, H3.3 ko, Daxx ko and Morc3 ko ES cells. Bar graph displays fold enrichment over a negative control region.

Next, to investigate if the Morc3 ATPase cycle would change this interaction context, we performed ChIP-MS experiments with the Morc3 mutant rescue cell lines. Here we focused on the ATP binding and the CW mutants which maintained proper localization with Morc3 targets (Supplementary Figure 5B,C) Comparing ChIP-MS results between wild type Morc3 and CW mutant Morc3 we did not detect striking differences (Figure 6C). In contrast, the ATP binding mutant resulted in distinct changes of the protein interaction network (Figure 6C). Focusing on the major ERV silencing factors, we found that they were still associated with ATP binding mutant Morc3. Interestingly, Daxx was specifically depleted in the ATP binding mutant suggesting that dimerization of Morc3 is critical to mediate Daxx interaction.

Daxx is known to regulate ERV silencing through a C-terminal SUMO interaction motif (SIM) (Sadic et al., 2015). Since several ERV silencing factors are SUMOylated, we set out to determine which proteins Daxx might interact with though this C-terminal SIM. We performed ChIP-MS experiments comparing Daxx knock-out cell lines with re-expression of wild type Daxx protein or Daxx^ΔSIM^. Lack of the C-terminal SIM reduced association with SUMO proteins and, importantly, resulted in strongly reduced binding with Morc3 (Figure 6D). Together, these data suggest that SUMOylated Morc3 interacts with Daxx through the C-terminal SIM.

Recent studies have shown the role of Daxx-mediated H3.3 incorporation in ERV silencing and suggest a dynamic turnover of histones in ERV heterochromatin (Elsasser et al., 2015; Navarro et al., 2020; Wolf et al., 2017). We therefore asked if association of Morc3 with Daxx is critical for H3.3 incorporation on Morc3 target sites. ChIP-seq profiles for H3.3 demonstrate that this histone variant is prominently enriched on Morc3 peaks (Figure 6E). We then performed ChIP-qPCR for H3.3 on selected Morc3 target regions in wild type, H3.3 ko, Daxx ko and Morc3 ko ES cells. Our data show that the H3.3 ChIP is specific, as signal is lost in H3.3 ko cells (Figure 6F). Consistent with the role of Daxx in H3.3 incorporation, H3.3 is strongly reduced in Daxx ko cells. Importantly, Morc3 ko cells also display strongly reduced H3.3 signal (Figure 6F), demonstrating that Morc3 is critical for Daxx-mediated H3.3 incorporation on ERVs.

## DISCUSSION

Our data show that Morc3 is a critical regulator of ERV silencing in mouse ES cells. Other members of the MORC protein family have already been implicated in ERV silencing. Morc1 was found to regulate transposable elements in the mouse germline (Pastor et al., 2014) and Morc2a regulates LINE1 repression in mouse ES cells (Fukuda et al., 2018). In human cells MORC2 was identified to influence HUSH complex silencing (Liu et al., 2017; Tchasovnikarova et al., 2017). Although the MORC family of proteins represent important regulators of ERV silencing in different cell types, their mechanism of action was largely obscure. We can show a specific function of Morc3 in regulating Daxx-mediated H3.3 incorporation to maintain ERV heterochromatin. We found that in absence of Morc3, ERV heterochromatin loses H3K9me3, shows increased accessibility and reduced levels of H3.3. A previous study has demonstrated high H3.3 turnover on ERV regions (Navarro et al., 2020). Smarcad1 evicts nucleosomes from ERV heterochromatin and, Daxx-dependent replacement by H3.3-containing nucleosomes is needed to maintain low chromatin accessibility. Loss of H3.3 or Daxx, therefore, leads to more accessible chromatin and less H3K9me3 on ERVs. We found, that Morc3 knock-out cells display a very comparable phenotype, supporting the notion that Morc3 function is needed for proper Daxx activity. Based on our data we propose the following model for this process. High nucleosome turnover on ERV heterochromatin is induced by chromatin remodelers, such as Smarcad1. Re-establishment of these nucleosomes is mediated by Daxx, which contributes H3.3/H4 dimers. For this activity, Daxx needs to interact with SUMOylated Morc3 in an ATP-bound state. ATP hydrolysis by Morc3 mediates ATPase dimer dissociation, a conformational change that could be necessary for Daxx release and may function as “licensing step” for Daxx-mediated H3.3 deposition. In agreement with this model, we found that (I) Daxx requires the C terminal SIM domain to interact with Morc3, (II) Morc3 is a highly SUMOylated protein (Mimura et al., 2010), (III) Daxx interaction is not detected with the dimerization-defective Morc3 ATP binding mutant, (IV) reduced ATP hydrolysis of the Morc3 CW mutant fails to re-establish ERV silencing. In this context it is interesting to note that proper ATP hydrolysis is inhibited by CW-ATPase domain interaction (Zhang et al., 2019b). Ligand binding by the CW domain relieves this inhibition and appears necessary for the Morc3 ATPase cycle. H3K4me3 was previously shown as ligand for the Morc3 CW domain, however, in ERV heterochromatin, we did not detect substantial levels of this modification. Therefore, we suspect that either H3K4me3 is present transiently or, other ligands for the CW domain exist that act in the context of ERV heterochromatin. An efficient ATPase cycle is not only critical for Morc3 function, but also affects the silencing activity of MORC2 (Douse et al., 2018). It will thus be interesting to see if the other members of the MORC family also regulate Daxx function in different genomic contexts and whether this function is critically linked with the different diseases associated with mutations in MORC family members (Hong et al., 2017; Jadhav et al., 2016; Sloan et al., 2016; Tchasovnikarova et al., 2017).

## MATERIAL AND METHODS

### Cell culture

Feeder-independent ES cells were cultured in ES cell medium (500 ml High Glucose DMEM (Sigma, D6429), 91 ml (15%) Fetal Bovine Serum (Sigma, F7524), 6.05 ml Penicillin-Streptomycin (Sigma, P4333), 6.05 ml MEM Non-Essential Amino Acid Solution (Sigma, M7145), 1.2 ml 0,35 % 2-Mercaptoethanol (Sigma M7522) and 2.4 ml homemade LIF) on 10 cm and 15 cm gelatin-coated plates.

### Lentivirus production

Lentiviral particles were produced to perform Daxx rescue, to carry reporter plasmids, to introduce single sgRNAs for CRISPR/Cas9 knock-out and to deliver the screening libraries. 293T cells were transfected with a mix of 24 μg plasmid DNA consisting of 8 μg lentiviral transfer vector, 8 μg of each of the packaging plasmids psPAX2 (#183) and pLP-eco-env (#811) mixed with 120 μl 2.5 M CaCl_2_ and adjusted to a final volume of 1200 μl with H_2_O. Subsequently 1200 μl 2xHBS solution (50 mM HEPES, 280 mM NaCl, 1.5 mM Na_2_HPO_4_, adjusted to pH 7.05 with NaOH) was added slowly and drop wise to the mix while vortexing. The transfection mix was added immediately to the 293T cells seeded one day ahead at a density of 4 million per 10 cm dish. Four to eight hours after transfection, the medium that contained precipitates of calcium phosphate and DNA was removed, cells were washed with 1xPBS and fresh medium was added. Virus containing culture supernatant was harvested 48 h after transfection.

### Lentiviral transduction of mouse ES cells by spinoculation

For lentiviral transduction 200.000-250.000 ES cells per well were seeded on gelatinized 6-well dishes in ES medium containing 8 μg/ml polybrene (Sigma, #H9268). Viral supernatant was added to the cells and plates were spun in a prewarmed centrifuge (Heraeus Multifuge 4KR, Thermo Electron Corporation) at 1000 x g for 1 h at 34°C to enhance viral transduction by improved viscosity for infection. After centrifugation the medium was carefully replaced or diluted with new ES medium to reduce toxic side effects of polybrene.

### sgRNA screen

The genome wide sgRNA screen was conducted using the 2-vector system (lentiGuide-Puro) GeCKOv2 pooled library. The plasmid library was obtained from the Zhang lab (Addgene catalog #1000000053) and amplified in E. coli as previously described (Sanjana et al., 2014). To ensure complexity of the library the purified plasmid pool was sequenced. Plasmids were used for lentivirus production by calcium phosphate transfection of 293T cells. To ensure complexity of the library ten parallel virus preparations with a total amount of 80 μg plasmid DNA were setup. Harvested supernatants were pooled. To identify optimal virus concentration for achieving a multiplicity of infection (MOI) of 0.3, virus titers were determined for each virus lot. Infectivity was tested by transducing ES cells with the GeCKOv2 library through spinoculation of 300.000 cells per well in 6-wells with different volumes of virus supernatant (between 1 and 100μL). After 48 hours, cells were transferred to 15cm plates and put under selection with 0.5 μg/ml puromycin until colonies were detected. The screening experiment was performed using T90 cells, which are ES cells carrying the TetO-EGFP-T2A-Zeo-GAG2.22 reporter at one specific integration (Sadic et al., 2015). T90 cells also carry the rtTA for doxycycline induction of the TRE promoter and a Cas9 transgene. For saturation, we aimed for infection of 100 cells per sgRNA. As the libraries carry 65.000 different sgRNAs, at least 6.5 million ES cells ought to be transduced. As the MOI should be below 30%, 30 million T90 cells were transduced with the lentiviral sgRNA library pool at a MOI between 27-28%. Transduced cells were selected by puromycin treatment after two days with 0.5 μg/ml puromycin. The SHIN reporter was activated with doxycycline two days after transduction with 0.1 μg/ml doxycycline. To determine the distribution of sgRNAs without selection pressure, the cell pool was dived in two groups. The control pool was cultured in medium containing 0.5 μg/ml puromycin and 0.1 μg/ml doxycycline to determine the baseline sgRNA distribution. The selection pool was treated with either 25 or 50 μg/ml zeocin from day 4 on. All cells were passaged every 2 days and harvested 8 days after transduction. For the control pool, 20 million cells and for the zeocin treated pool 400.000 cells where collected. Extraction of genomic DNA was performed using the DNeasy Blood and Tissue Kit Mini (Qiagen, #69504). DNA was sheared using a syringe and 27 g needle and DNA concentration was determined with the Q-bit fluorometer (Invitrogen). Quantitative PCR (qPCR) was used on genomic DNA extracted from ES cells in the library screen to determine the number of cycles necessary for amplification of the sgRNA sequences with primers GS3369 and GS3371. PCR was carried out with the Fast SYBR® Green Master Mix™ (Applied Biosystems) in a LightCycler480™ (Roche). The reactions were performed in a total volume of 20 μl in a 384 well plate (Sarstedt). Ct-values were generated by the LightCycler480-Software (Roche) using the 2nd derivative max function. The library preparation was performed in two steps of PCR reactions: For the first PCR, the amount of input genomic DNA (gDNA) for each sample was calculated to achieve 100x coverage over the GECKO library, which resulted in an input of at least 40 μg DNA per sample (assuming 6.6 μg of gDNA for 1 million cells). For the control sample, 46 separate 60 μl PCR reactions were performed with 1 μg gDNA in each reaction using Q5® High-Fidelity DNA Polymerase (New England Biolabs). For the zeocin treated sample the total amount of genomic DNA harvested was used. Primers used to amplify lenti CRISPR sgRNAs for the first PCR were: GS3367 and GS3368. The first PCRs were pooled and 3 μl were used as template for the second PCR. The second PCR served to attach Illumina adaptors and to barcode samples and was done in a 60 μl reaction volume divided to two times 30μl. Primers for the second PCR include both a variable length sequence to increase library complexity and an 8bp barcode for multiplexing of different biological samples Amplification was carried out with 18 cycles for the first PCR and 30 cycles for the second PCR. Primers for the second PCR for zeocin treated samples were GS3369 and GS3371 (Index9) and for the control samples GS3369 and GS3370 (Index11). Resulting amplicons of the second PCR were mixed, purified using AMPure XP beads (Beckman Coulter) in a volume ratio of 1:1 according to manufacturer’s instructions. Prior to sequencing, purified PCR product was quality controlled by Qubit fluorometer (Invitrogen) and Bioanalyzer (Agilent Technologies) measurement according to manufacturer’s instructions. Sequencing was carried out by LAFUGA sequencing facility at the Gene Center of the LMU Munich using the Illumina HiSeq 1500 and a 50bp single end run.

### SHIN silencing assays

#### Initiation assay

SHIN Reporter constructs (#940 EGFP only and #1074 EGFP SHIN) were stably integrated into cells by lentiviral transduction and the percentage of EGFP^+^ cells was measured by FACS after 2-4 days. Lentiviral particles were generated using standard protocols and virus titers were determined by titration in T37 cells. Mouse ES cells were transduced on gelatinized multi-well dishes using spinoculation at a low multiplicity of infection to ensure a linear relationship between virus titer and transduction rate. The ratio of the percentage of EGFP^+^ cells generated by the reporter relative to the percentage of EGFP^+^ cells generated by a control EGFP vector of the same virus titer was used to quantify reporter silencing (relative %EGFP+ cells).

#### Maintenance assay

Cells containing the SHIN reporter (based on T90 cells) were incubated with 0.1 μg/ml doxycycline for 2 or 4 days and expression of the reporter locus was measured as percentage of EGFP^+^ cells. To test the effect of potential ERV silencing mutants, T90 cells were transduced with lentiviral sgRNA constructs. Two days after transduction, cells were selected for sgRNA expression with 1 μg/ml puromycin and SHIN reporter was induced with doxycycline.

### Morc3 FLAG knock-in

The Morc3-3xFLAG knock in cell line K14-E8 was generated by CRISPR mediated double strand break induction close to the Morc3 STOP codon followed by homologous recombination providing a template of the same genomic region (#1529, Supplementary Table S8), including the 3xFLAG tag before the STOP and exchanging the sequence of the PAM from TGG to TAG so that the repaired sequence cannot be targeted (Supplementary Figure S1). CRISPR and homology plasmids were transfected via jetPRIME (Polyplus-transfection) and single clones were analyzed for Morc3-3xFLAG expression by western blot.

### CRISPR /Cas9 mediated knock-out

Stable knock out cell lines were generated by small guide RNA (sgRNA) mediated Cas9 DNA cleavage using the pX330 plasmid (Cong et al., 2013; Hsu et al., 2013, Addgene plasmid #42230). DNA oligonucleotides were hybridized and ligated into the BbsI digested pX330 to introduce the sgRNA sequence into the vector. Mouse ES cells were co-transfected with the pX330 plasmid harboring the sgRNA and a plasmid encoding a puromycin resistance gene (pLFIP) using jetPRIME (Polyplus-transfection). After two days,transfected cells were selected by addition of 2 μg/ml puromycin to the medium. Puromycin selection was removed after 1 day, and individual cell clones were isolated after 4-6 days. Clonal cell lines were analyzed by western blotting or by PCR and Sanger sequencing.

### Rescue cell lines

For generation of Morc3 rescue cell lines, KO27-2 cells were transfected with the PiggyBac (PB) Transposon Vector System via jetPRIME (Polyplus-transfection). The system consists of two plasmids, one encoding the transposase (#1704) and a second one containing the respective rescue construct (Supplementary Table S7). After two days, transfected cells were selected by addition of 2 μg/ml puromycin to the medium. Individual cell clones were isolated while puromycin selection was maintained for 7-10 days.

For Daxx rescue cell lines, the Daxx KO cell line KO2-3 was transduced with lentivirus carrying the plasmids #1272 or #1273, respectively (Supplementary Table S7).

### RT-qPCR

Total RNA was extracted using Trizol and the RNA Clean & Concentrator −25 Kit (Zymo research, #R1017) including on-column DNAse digestion (Qiagen, #79254) according to manufacturer’s instructions. For cDNA synthesis, 1 μg of RNA was used as input. The reaction was carried out using SuperScript III reverse transcriptase (Invitrogen, #18080044), Random Primer 6 (NEB, #S1230S), RNasin Ribonuclease inhibitor (Promega, #N2515). First, total RNA, random hexamer primers and RNase-free water were mixed and incubated for 10 min at 70°C, followed by one-minute incubation on ice. Next, SuperScript buffer, dNTP and DTT were added. Additionally, rRNasin and SuperScript III reverse transcriptase were added and incubated at 25°C for 8 min, followed by incubation at 50°C for 50 min. Reaction was stopped by heat inactivation at 70°C for 15 min. qPCR was carried out with the Fast SYBR® Green Master Mix (Applied Biosystems, 4385612) in a LightCycler480™ (Roche) according to the Fast SYBR Green Master Mix-protocol. Primers were evaluated for generating a single PCR product and for linear amplification in a wide range of DNA template dilutions. Every PCR-reaction was performed in a total volume of 10 μl in triplicates in a 384-well plate (Sarstedt). Two independent control genes (Actin and HPRT) were used as reference genes for RT-qPCR experiments and geometric mean of reference Ct values was used as normalization as described (Vandesompele et al., 2002). Ct-values were generated by the LightCycler480-Software (Roche) using the 2nd derivative max function and fold changes were calculated using the 2^−ΔΔC^ method.

### ChIP-seq

For the standard ChIP protocol, 25 million ES cells were cross-linked in 7-8 ml pre-tempered ES medium containing 1% formaldehyde (Pierce #28906) for 10 min at 22°C. Fixation was stopped by addition of 0.125 M final concentration of glycine, followed by two washing steps with PBS containing 10% Serum. Fixed cells were resuspended in 10 ml ice cold buffer LB1 (50 mM Hepes-KOH pH 7.5, 140 mM NaCl, 1 mM EDTA, 10% Glycerol, 0.50% NP-40, 0.25% Triton X-100, 1x Roche cOmplete Mini, EDTA-free Protease Inhibitor Cocktail (Roche, #04693159001), rocked at 4°C for 10 min and after centrifugation resuspended in 10 ml ice cold buffer LB2 (10 mM Tris-HCl pH 8.0, 200 mM NaCl, 1 mM EDTA, 0.5 mM EGTA, 1x Roche cOmplete protease inhibitors), rocked at 4°C for 5 min and centrifuged again. The pelleted nuclei were resuspended in 1 ml ice cold shearing buffer (10 mM Tris-HCl pH 8.0, 100 mM NaCl, 1 mM EDTA, 0.5 mM EGTA, 0.1% Na-Deoxycholate, 0.1% SDS, 1x Roche cOmplete protease inhibitors). Buffer compositions for LB1, LB2 and shearing buffer are based on (Boyer et al., 2005). Exactly 1 ml nuclei suspension was transferred to a 1 ml milliTUBE with AFA fiber (Covaris, #520130) and sonicated using a Covaris S220 device for a time of 15-20 min and the following settings: Peak power 140 Watt, Duty factor 20%, Cycles per burst 200, Temperature 4°C. The sheared samples were added to a 1.5 ml tube with 110 μl of 10% Triton X-100 (final concentration 1%) and centrifuged at 14000 rpm for 15 minutes at 4°C. The soluble chromatin containing supernatant was divided in 110 μl aliquots (equivalent to 2-2.5 mio cells) and for IP diluted with 890 μl complete Buffer A (10 mM Tris-HCl pH 7.5, 1 mM EDTA, 0.5 mM EGTA, 1% Triton X-100, 0.1% SDS, 0.1% Na-Deoxycholate, 140 mM NaCl, 1x Roche cOmplete protease inhibitors). Per IP 30 μl of Protein G Dynabeads (Invitrogen, #10004D) were used. For FLAG ChIP-seq experiments 1-2 μl of FLAG M2 antibody (Sigma-Aldrich, #F3165) were used per IP and 4-5 IPs (equivalent to 10-12 million cells) were pooled to obtain enough material for library preparation. For H3K9me3 ChIP-seq experiments 1 μl of H3K9me3 antibody (Active Motif, #39161) was used per IP. Beads were incubated with the corresponding antibody in complete Buffer A for 1.5 h prior to IP, followed by two washes with Buffer A (without Roche cOmplete protease inhibitors). Diluted Chromatin was added to the pre-bound beads and incubated for 4 h on a rotating wheel with 30 rpm at 4°C. After IP beads were washed 5 times with Buffer A and 1 time with Buffer C (10 mM Tris-HCl pH 8.0, 10 mM EDTA) beads were resuspended in 100 μl elution buffer (10 mM Tris HCl pH 8.0, 300 mM NaCl, 5 mM EDTA, 0.5% SDS). RNA was degraded with 2 μl RNase A (10 mg/ml) for 30min at 37°C. Proteins were digested with 2 μl Proteinase K (10 mg/ml) at 55°C for 1 h and crosslink reversal of immunoprecipitated DNA was carried out overnight at 65°C. DNA was purified using the Agencourt AMPure XP beads (Beckman Coulter, #A63882).

For H3K9me3 ChIP-seq in wild type and Morc3 ko cells, a protocol was adapted from (Dahl and Collas, 2008). Briefly, 2 mio cells were lysed in 100 μl Buffer-B (50 mM Tris-HCl, pH 8.0, 10 mM EDTA, 1% SDS, 1x Roche cOmplete protease inhibitors) and sonicated in a microtube (Covaris; #520045) using the Covaris S220 device with the settings: Peak power 105 Watt, duty factor 2%, cycles per burst 200, Temperature 4°C. The supernatant after centrifugation was diluted with 900 μl complete Buffer A and 150 μl chromatin (corresponding to approximately 300.000 cells) were used per IP with 10 μl Protein G Dynabeads and 1 μl of H3K9me3 antibody as described above.

Library preparation was performed using the Ultra II DNA Library prep kit for Illumina (NEB, #E7645S) according to manufacturer instructions. Sequencing was performed by LAFUGA on an Illumina Hiseq 1500 using 50 bp paired end run.

### ATAC-seq

The OmniATAC transposition reaction was performed with 50000 mES cells as previously described (Corces et al., 2017) using the Tagment DNA TDE1 Enzyme (Illumina, #20034197). DNA was purified using the PCR clean-up MinElute kit (Qiagen, #28006). The transposed DNA was subsequently amplified in 50μl reactions with custom primers as previously described (Buenrostro et al., 2013). Libraries were purified and size selected for fragments less than 600bp using the Agencourt AMPure XP beads (Beckman Coulter, #A63882). Sequencing was performed by LAFUGA on the Illumina Hiseq 1500 with 50 bp single end reads.

### RNA-seq

For samples GS271-GS276, total RNA was extracted using RNeasy Mini Kit (Qiagen, #74106) including on-column DNAse digestion (Qiagen, #79254). Ribosomal RNA was depleted using RNA Ribo-Zero rRNA Removal Kit (Illumina, #MRZH11124). For sample GS948, total RNA was extracted using Trizol and the RNA Clean & Concentrator −25 Kit (Zymo research, #R1017) including on-column DNAse digestion (Qiagen, #79254). Due to the discontinued Ribo-Zero kit, ribosomal RNA was depleted with the NEBNext rRNA Depletion Kit (NEB, #E6350). Libraries were prepared with the NEBNext Ultra Directional RNA Library Prep Kit for Illumina (NEB, #E7420S). Sequencing was performed by LAFUGA on an Illumina Hiseq 1500 using 50 bp paired end run.

### ChIP-MS

Immunoprecipitation of bait proteins (Daxx or Morc3) was performed according to the “Rapid immunoprecipitation mass spectrometry of endogenous protein (RIME) for analysis of chromatin complexes” protocol (Mohammed et al., 2016). Cells were harvested and fixed at 22°C for 10 minutes in ES medium with 1% Formaldehyde (Pierce #28906) with a density of 4×10^6^ cells/ml. The fixation was stopped by 5 min incubation with glycine at a final concentration of 0.125 M. The cell pellet was washed twice with ice cold PBS containing 10% serum and once with ice cold PBS without serum. Cell pellets of 60×10^6^ cell were flash frozen and stored in −80°C for later use. All following steps were performed in 4°C or on ice and centrifugation (Thermo scientific, Heraeus multifuge X3R) was performed at 2000g for 5 minutes if not specified. For each immunoprecipitation experiment 60×10^6^ cells and 60 μl of Protein G Dynabeads (Invitrogen, #10004D) coupled with 6 μl of the FLAG M2 antibody (Sigma-Aldrich, #F3165) were used. Beads were washed twice with LB3 (10 mM Tris-HCl pH 8.0, 100 mM NaCl, 1 mM EDTA, 0.5 mM EGTA, 0.1% Na-Deoxycholate, 0.5% N-Lauroylsarcosine) and resuspended in 100 μl complete LB3 (LB3 containing 1x Roche cOmplete Mini, EDTA-free Protease Inhibitor Cocktail (Roche, #04693159001)). The antibody was prebound to beads at 35 rpm on a rotating wheel for 2-4 h and the beads were washed 4×1 mL with LB3.

Cells were lysed by resuspending in complete LB1 (50 mM Hepes-KOH, pH 7.5, 140 mM NaCl, 1 mM EDTA, 10% glycerol, 0.5% NP-40, 0.25% Triton X-100, 1x Roche cOmplete Mini, EDTA-free Protease Inhibitor Cocktail (Roche, #04693159001)) and incubation at 35 rpm for 10 min followed by centrifugation. To obtain nuclei, the lysed cells were resuspended in LB2 (10 mM Tris-HCl, pH 8.0, 200 mM NaCl, 1 mM EDTA, 0.5 mM EGTA, Roche cOmplete protease inhibitors) and incubated at 35 rpm for 5 min followed by centrifugation. The nuclei were resuspended in complete LB3 (complete LB3, 1x PhosStop EASYpack (Roche, #4906837001), 20 mM N-Ethylmaleimide (NEM) (Thermo Fisher, #23030)) at a maximum density of 20 mio cells per 300 μl. Chromatin was sheared with Bioruptor pico (Diagenode) for 10-12 cycles with 30 sec on and 30 sec off. After shearing a 1/10 volume of Triton X-100 was added followed by centrifugation at 20.000g for 10 min. The supernatant was added to the beads for chromatin-immunoprecipitation and incubated for 4 h at 35 rpm on a rotating wheel. The beads with the bound chromatin were washed with 4×1 ml of RIPA buffer (50 mM Hepes-KOH pH 7.6, 500 mM LiCl, 1 mM EDTA, 1% NP-40, 0.7% Na-Deoxycholate) for 5 min at 20 rpm on a rotating wheel followed by the same washing process with 2×1 ml freshly prepared ice cold 100 mM AMBIC (ammonium hydrogen carbonate) solution. On the last AMBIC wash the beads were transferred to a new tube. The beads were stored dry at −80°C until MS analysis.

The sonicated chromatin was size checked with a 2100 Bioanalyzer (Agilent) using the DNA high sensitivity kit (Agilent, #5067-4626) or the DNA 7500 kit (Agilent, #5067-1506) following manufactures protocol. Shortly, Elution buffer (0.5% SDS, 300 mM NaCl, 5 mM EDTA, 10 mM Tris HCl) was added 1:10 to the sonicated chromatin and incubated at 65°C, 800 rpm overnight for de-crosslinking. The de-crosslinked chromatin was treated with 0.2 μg/ml of RNAse A and incubated at 37°C, 700 rpm for 1 h. Next, the sample was treated with 0.2 μg/ml Proteinase K and incubated at 55°C, 700 rpm for 2 h. The DNA was purified with the MinElute PCR purification kit (Qiagen, #28006) and concentration was measured with the Qubit dsDNA high sensitivity kit (Thermo Fisher, #Q32851) before loading the samples to the DNA HS/DNA 7500 chip.

### Liquid chromatography – mass spectrometry (LC–MS)

Beads were washed three times with 50 mM NH_4_HCO_3_ and incubated with 0.5 μg/μl Lys-C and 20 U benzonase in 6 M urea 50 mM NH_4_HCO_3_ pH 7.5 for 90 min at 28°C, washed with 50 mM NH_4_HCO_3_ and the combined supernatants were digested overnight with 0.2 μg/μl of trypsin in presence of 10 mM DTT. Digested peptides were alkylated with 30 mM IAA and desalted prior to LC-MS analysis.

For LC-MS/MS purposes, desalted peptides were injected in an Ultimate 3000 RSLCnano system (Thermo), separated in either a 15 cm analytical column (75 μm ID with ReproSil-Pur C18-AQ 2.4 μm from Dr. Maisch) with a 50 min gradient from 4 to 40% acetonitrile in 0.1% formic acid or in a 25 cm analytical column (75 μm ID, 1.6 μm C18, Aurora-IonOpticks) with a 50 min gradient from 2 to 35% acetonitrile in 0.1% formic acid. The effluent from the HPLC was directly electrosprayed into a Qexactive HF (Thermo) operated in data dependent mode to automatically switch between full scan MS and MS/MS acquisition. Survey full scan MS spectra (from m/z 375–1600) were acquired with resolution R=60,000 at m/z 400 (AGC target of 3×10^6^). The 10 most intense peptide ions with charge states between 2 and 5 were sequentially isolated to a target value of 1×10^5^ and fragmented at 27% normalized collision energy. Typical mass spectrometric conditions were: spray voltage, 1.5 kV; no sheath and auxiliary gas flow; heated capillary temperature, 250°C; ion selection threshold, 33.000 counts.

MaxQuant 1.6.14.0 was used to identify proteins and quantify by iBAQ with the following parameters: Database, UP000000589_10090_Mmusculus_2020; MS tol, 10 ppm; MS/MS tol, 20 ppm Da; Peptide FDR, 0.1; Protein FDR, 0.01 Min. peptide Length, 7; Variable modifications, Oxidation (M); Fixed modifications, Carbamidomethyl (C); Peptides for protein quantitation, razor and unique; Min. peptides, 1; Min. ratio count, 2.

### Immunofluorescence analysis

Cells were seeded in 24 well plates containing 12mm poly-L-lysine coated coverslips (50.000 cells / well). Cells were fixed with formaldehyde (500 μl/well, 3.7% formaldehyde in PBS for 10 min at RT). Coverslips were washed twice for 5 min in 1 ml PBS. Permeabilization was performed for 5 min in 500 μl/well in permeabilization solution (10 mM sodium citrate tribasic dihydrate, 200 μl Triton X-100). Two washing steps with 1 ml PBS followed by two washing steps with 1 ml washing solution I (2 l H_2_O, 220 ml 10 x PBS, 2.2 ml Tween-20, 5.5 g BSA) were performed before samples were blocked for 30 min in 300 ml blocking solution I (50 ml washing solution I, 1.2 g BSA). Primary antibody was diluted in 200 μl/well blocking solution I and incubated over night at 4°C in the dark. Cells were then washed 3 times with washing solution I. Secondary antibody was diluted in 200 μl/well blocking solution II (0.5 ml blocking solution I, 0.5 ml serum (goat or donkey)) and incubated for 1 h at RT in the dark. Next, coverslips were washed 3 times 10 min with washing solution II (500 ml PBS, 500 μl Tween-20), followed by embedding coverslips with Vectashield containing DAPI (Vector Laboratories, Axxora - H-1200) and sealed with nail polish. Samples were stored at 4°C in the dark until pictures were taken at inverted confocal microscope Leica SP5 (64x) and analyzed with ImageJ.

### Bioinformatics analysis

#### sgRNA screen data analysis

Fastq files were trimmed with Trimmomatic version 0.36 and options “CROP:43 HEADCROP:23” to contain only the unique sgRNA sequence. Reads were then mapped with bowtie to all sgRNA sequences of the Gecko V2 library. Cvs files of library sequences were downloaded from addgene (https://www.addgene.org/pooled-library/zhang-mouse-gecko-v2/). CSV files were converted to fasta format using R. Bowtie was used to generate the library index. Reads were then aligned with bowtie 1. The number of aligned reads to each sgRNA sequence was calculated with bedtools command “bedtools genomecov”. The numbers of reads for each sgRNA per sample were normalized in R as follows: Normalized reads per sgRNA = (reads per sgRNA / total reads for all sgRNAs in sample) + 1. Hits were identified by conversion of sgRNA enrichment scores into gene rankings by statistical analysis using the SecondBestRank scoring method for RNAi gene enrichment ranking (RIGER) with the RigerJ tool.

#### ChIP-seq

Paired end ChIP-seq reads were aligned to the mouse genome mm10 using Bowtie2 with default settings. The resulting BAM files were filtered to remove non-paired reads, low mapping quality, non-primary alignment and PCR duplicates with “samtools view -b -f 2 -F 1280 -q 20”. Homer tag directories were generated with “makeTagDirectory”. BigWig files were generated from tag directories with “makeBigWig.pl mm10 -webdir . -url . -norm 1e7 -normLength 100 -fragLength 150 -update”.

Morc3 peaks were identified using findPeaks with option “-style factor” over Input. The peak files of replicate experiments were merged using homer “mergePeaks” and, peaks common in two replicates were kept for the final peak list. The final Morc3 peak list was annotated using homer “annotatePeaks.pl”, resulting in detailed peak annotation including association with repeat elements.

Morc3 association with repeats was determined using RepEnrich2. The fraction counts for each repeat family were normalized and repeat classes with significant enrichment (padj < 0.05 & log2FoldChange > 0.5) in Morc3-FLAG ChIP-seq vs. wild type FLAG ChIP-seq (background) were calculated using DeSeq2. Enrichment of H3K9me3 on ERVs in wild type vs. Morc3 knock-out and Morc3 rescue cells was also calculated with RepEnrich2 and DeSeq2.

Cumulative coverage plots on IAPEz elements was performed as described earlier (Sadic et al., 2015).

Heatmaps for ERV silencing factors Trim28, Setdb1, Morc3 and histone H3.3 / H3K9me3 were plotted as log-transformed normalized coverage (calculated from Homer tag directories with annotatePeaks.pl).

#### RNA-seq

Paired end reads were aligned to the mouse genome version mm10 using STAR with default options “--runThreadN 32 --quantMode TranscriptomeSAM GeneCounts --outSAMtype BAM SortedByCoordinate”. Read counts for all genes were normalized using DESeq2. Differentially expressed genes were determined using the DESeq2 results function (adjusted p-value <0.01).

Expression of ERV families was calculated using RepEnrich2. Fraction counts were normalized and differentially expressed repeats were calculated using DeSeq2.

#### ATAC-seq

Single end ATAC-seq reads were aligned to the mouse genome mm10 using Bowtie with options “-q -n 2 --best --chunkmbs 2000 -p 32 -S -m 1”. Duplicated reads were subsequently removed using Picard. Homer tag directories were generated using makeTagDirectory and bigwig files were generated with makeBigWig.pl. ATAC peaks were identified using Homer findPeaks.pl with option “-style factor” over Input. Peaks from all samples were merged using mergePeaks resulting in a unified Peak set. Raw ATAC coverage counts were calculated with annotatePeaks. Differential ATAC peaks were determined with the DESeq2 result function.

ATAC coverage of ERV families was calculated using RepEnrich2 on BAM files including multimapping reads. Fraction counts were normalized and differentially accessible repeats were calculated using DeSeq2.

#### IGV Screenshots

IGV screenshots were taken from respective bigwig files using IGV 2.5.0.

#### ChIP-MS data analysis

For data processing MaxQuant (1.16.14.0) was used together with a reference mouse protein database downloaded from UniProt (Proteome ID: UP000000589). The final list of proteins found was filtered and statistically processed in R studio version 3.5.0 using the Linear Models for Microarray Data (LIMMA) R script Version 1.0.1 that is available on GitHub written by Wasim Aftab (https://github.com/wasimaftab/LIMMA-pipeline-proteomics).

For comparative analysis between Morc3 wild type and mutant proteins, iBAQ values for proteins identified with more than 2 peptides were normalized to Morc3 or Daxx, respectively, by dividing by iBAQ value of the bait. Mean normalized iBAQ values from replicate experiments were plotted. Daxx was not detected in the Morc3 ATP binding mutant ChIP-MS experiment.

Pathway enrichment analysis of significantly Morc3-associated proteins was performed using Panther.

## DATA ACCESS

ATAC-seq, ChIP-seq and RNA-seq datasets were deposited to GEO.

## ACKNOWLEDGEMENTS

High throughput sequencing was performed by the Laboratory for Functional Genome Analysis (LAFUGA) of the Ludwig-Maximilian-University, Munich.

We acknowledge the Core Facility Flow Cytometry and the Bioinformatics Core Unit at the Biomedical Center, Ludwig-Maximilian-Universität München, for providing equipment, service and expertise.

Mouse GeCKOv2 CRISPR knockout pooled library was a gift from Feng Zhang (Addgene #1000000052).

## AUTHOR CONTRIBUTIONS

Conceptualization: S.G., G.S.

Investigation: S.G., A.V.M., L.M., C.V.S., H.B., G.P.dA., A.S., I.F.

Data Curation: S.G., A.V.M., L.M., C.V.S., H.B., G.P.dA., A.S., I.F.

Formal Analysis: S.G., A.V.M., A.S., I.F., G.S.

Funding Acquisition: G.S.

Methodology: S.G., G.S.

Project Administration: S.G., G.S.

Resources: A.I., G.S.

Software: S.G., A.S., I.F., G.S.

Supervision: S.G., G.S.

Validation: S.G., A.V.M., L.M., C.V.S., H.B., G.S.

Visualization: S.G., H.B., C.S., A.V.M., G.S.

Writing – Original Draft: S.G., G.S.

Writing – Review & Editing: S.G., A.V.M., L.M., C.V.S., H.B., G.P.dA., A.S., I.F., A.I., G.S.

---

**Figure S1.**
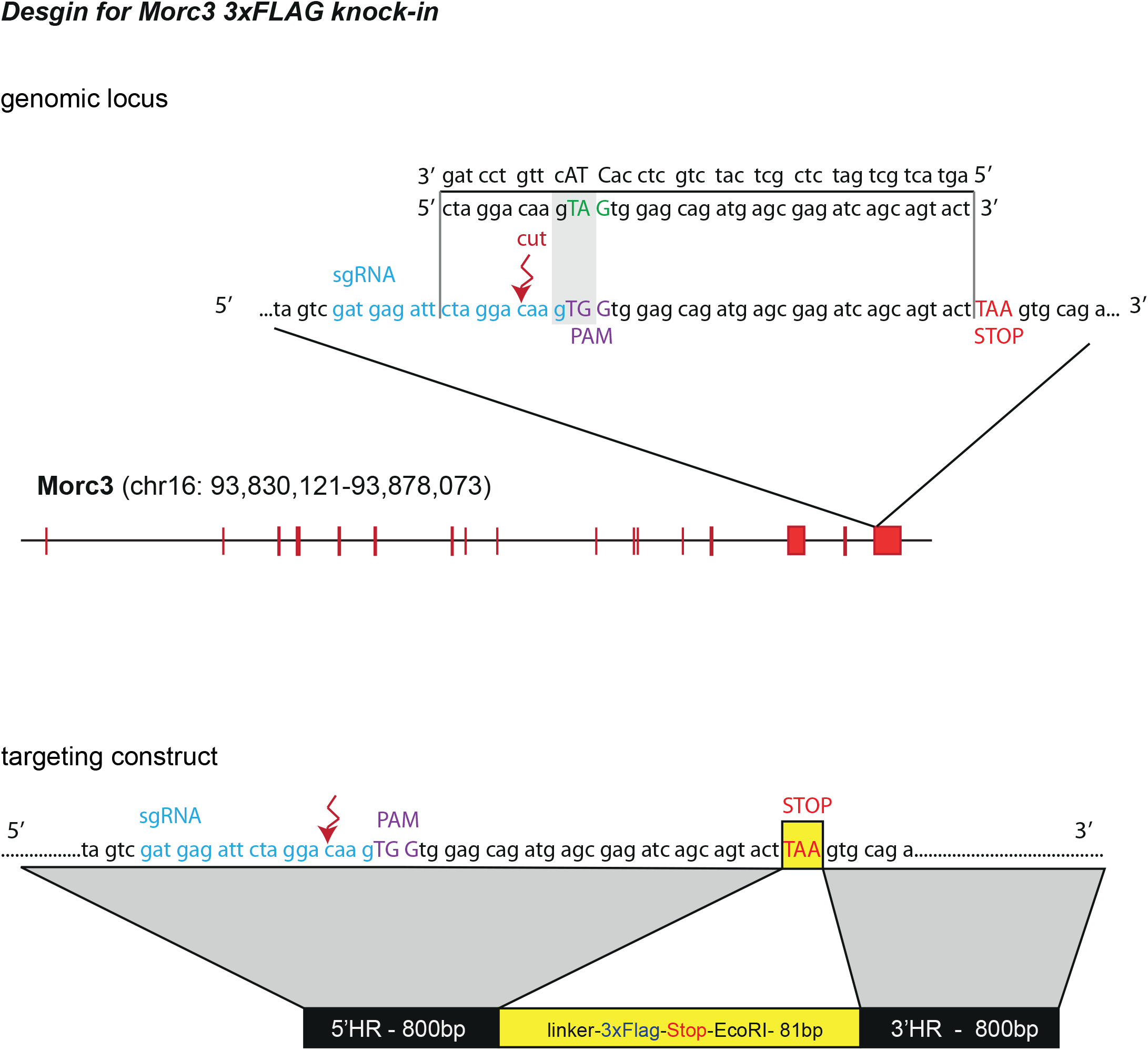
Morc3-3xFLAG knock-in strategy. Schema of the Morc3 genomic locus with the sequence that was used for sgRNA-mediated double strand break induction to allow for homology-dependent repair with the targeting construct.

**Figure S2.**
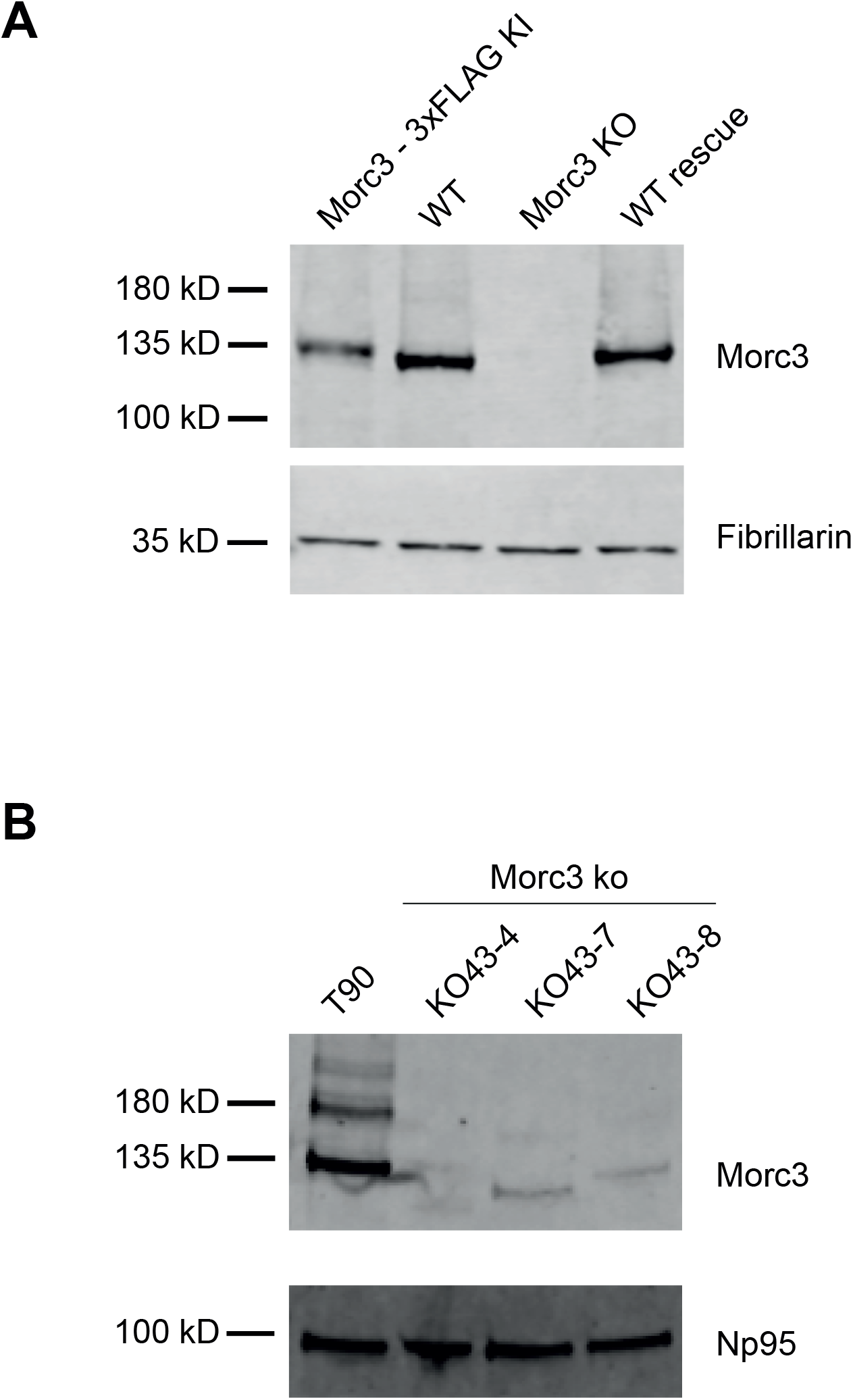
Validation of Morc3 knock-in, knock-out and rescue cell lines. (A) Western blot analysis of Morc3-3xFLAG knock-in, wild type, Morc3 ko and Morc3 rescue cell lines using antibodies against Morc3 and Fibrillarin (loading control). (B) Western blot analysis of Morc3 knock-out clones, based on T90 ES cells. Morc3 antibody staining indicates loss of Morc3 expression in the knock-out clones. NP95 western serves as loading control.

**Figure S3.**
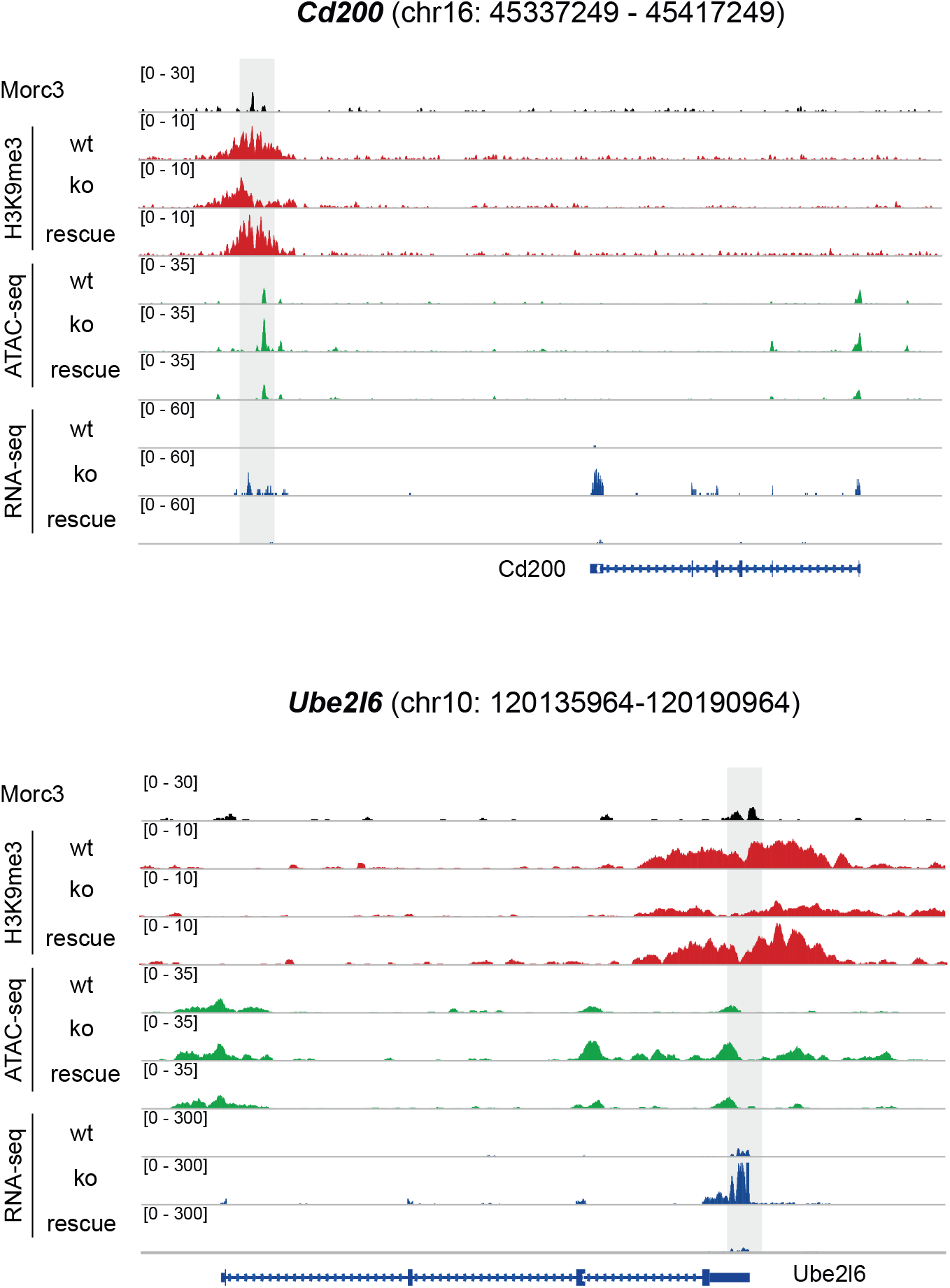
Morc3 dependent chromatin changes on selected Morc3 targets. Genome browser view of Morc3-dependent chromatin changes on target genes (Cd200, Ube2l6). Positions of Morc3 peaks are indicated by gray boxes. Chromatin and transcriptional changes are rescued in Morc3 rescue ES cells.

**Figure S4.**
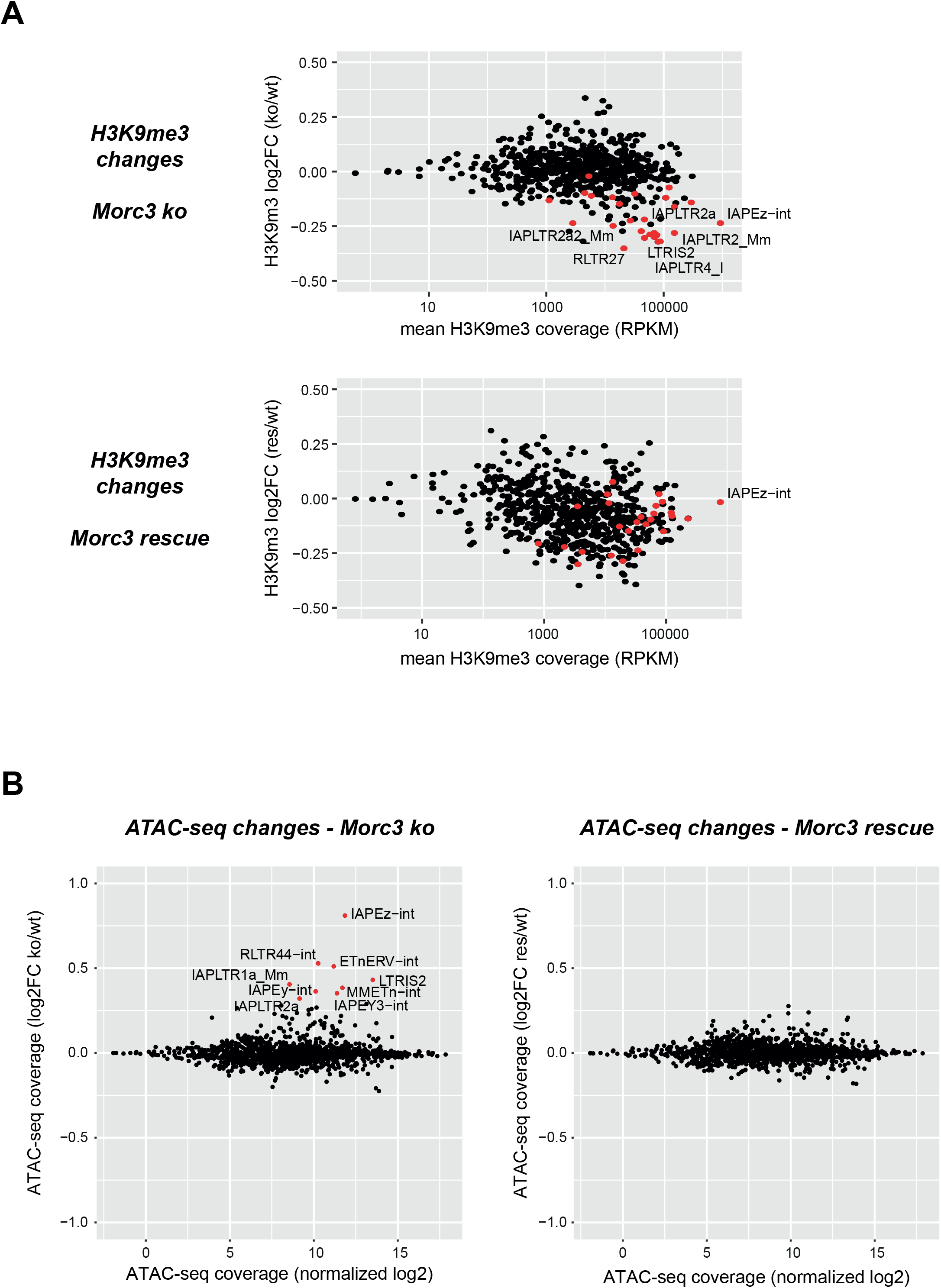
Chromatin alterations in Morc3 knock-out cells can be rescued. (A) Dot plot showing normalized H3K9me3 ChIP-seq coverage vs. log2-fold change of ERV families in wild type vs. Morc3 knock-out ES cells (upper panel) and wild type vs. Morc3 rescue ES cells (lower panel). Red dots indicate ERV families with significantly reduced coverage in Morc3 ko cells (adjusted p-value < 0.05, n=2 for each condition). H3K9me3 of most families is normalized in Morc3 rescue cells (e.g. IAPEz). (B) Dot plot showing average ATAC-seq coverage vs. log2-fold change of ERV families in wild type vs. Morc3 knock-out ES cells (left panel) and wild type vs Morc3 rescue ES cells (right panel. Colored dots indicate ERV families with significantly increased (red dots) or decreased (blue dots) coverage (adjusted p-value < 0.05, n=3 for each condition).

**Figure S5.**
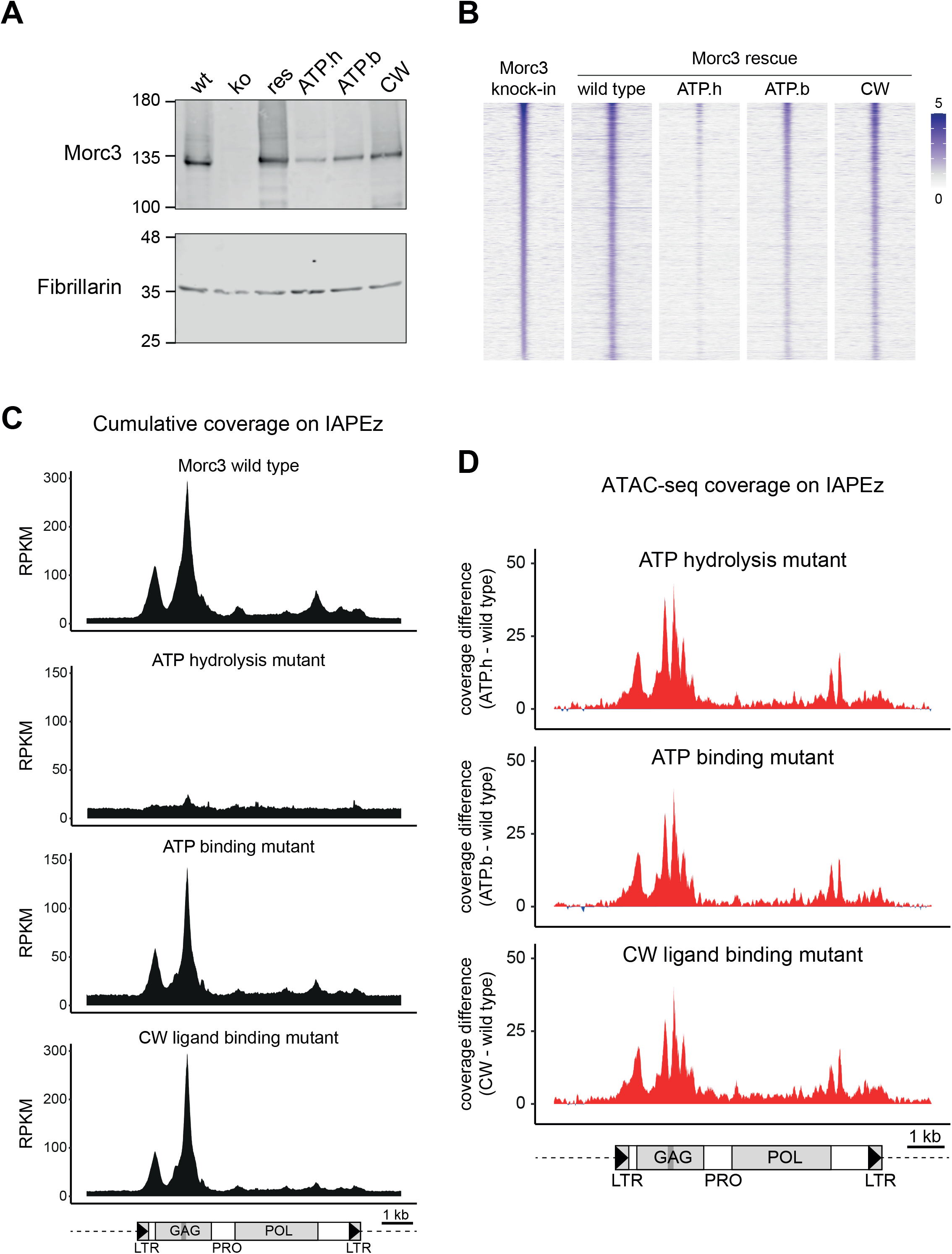
Morc3 ATPase mutants fail to rescue Morc3-dependent chromatin changes. (A) Western blot analysis of wild type, Morc3 ko and Morc3 rescue cell lines using antibodies against Morc3 and Fibrillarin (loading control). (B) Morc3 mutant proteins are recruited to wild type Morc3 peaks. Read-density heat map showing the normalized coverage of Morc3 wild type and mutant proteins on Morc3 binding sites. The Morc3 ATP hydrolysis mutant displays strongly reduced coverage on Morc3 peaks. (C) Cumulative coverage plot of Morc3 wild type and mutant proteins on IAPEz elements. Prominent enrichment is over the 5’UTR and the GAG region. The position of the SHIN sequence is indicated as dark gray bar. The ATP hydrolysis mutant does not display significant coverage on IAPEz elements. (D) Increased chromatin accessibility on IAPEz elements cannot be rescued by Morc3 mutant proteins. Plots display the difference in cumulative ATAC-seq coverage between Morc3 mutant and wild type ES cells. The position of the SHIN sequence is indicated as gray bar.

**Figure S6.**
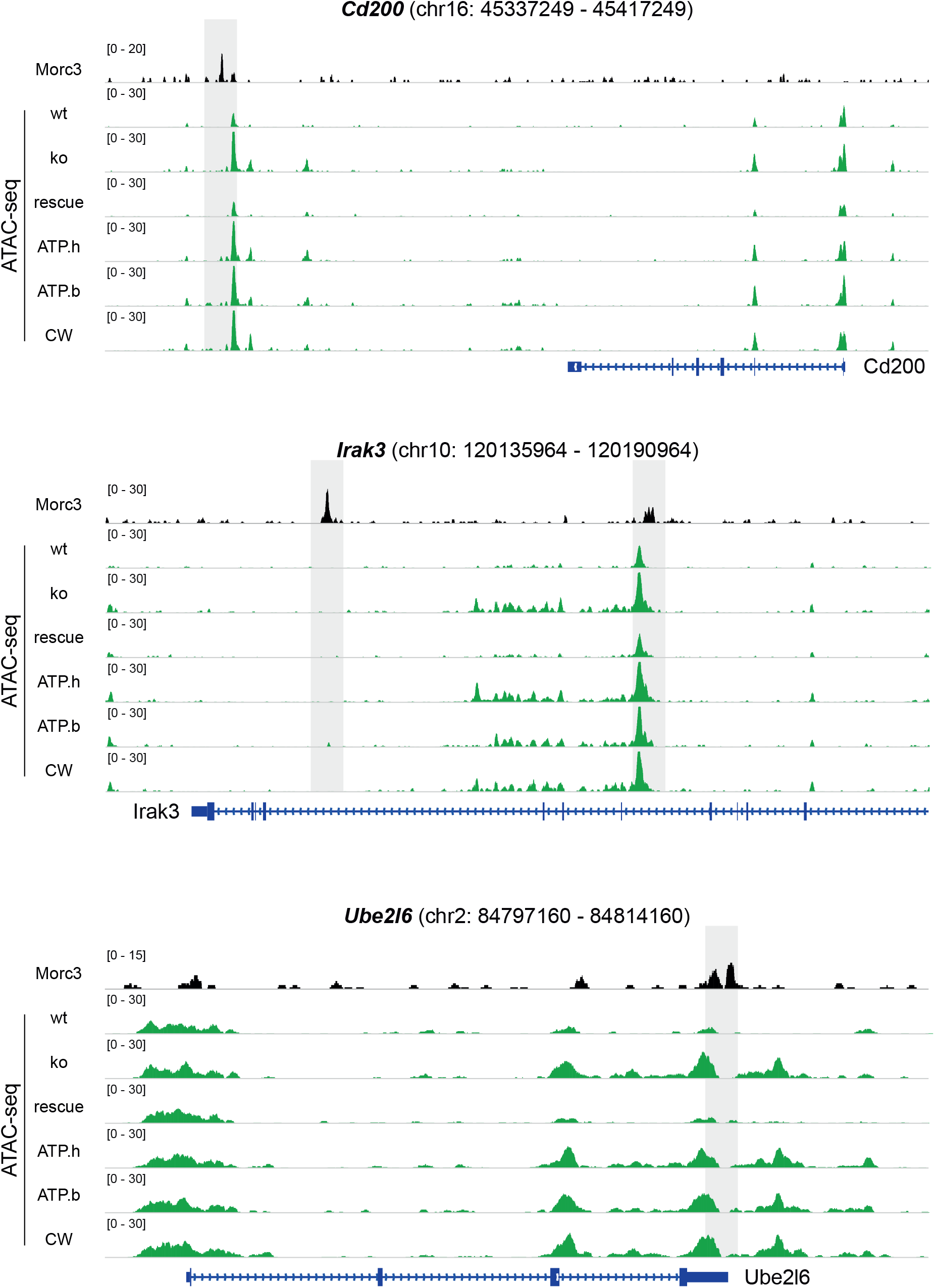
Morc3 mutant proteins fail to rescue increased chromatin accessibility on Morc3 targets. Genome browser view of chromatin accessibility on Morc3 target genes (Cd200, Irak3, Ube2l6). Positions of Morc3 peaks are indicated by gray boxes. wt – wild type, ko – Morc3 ko, rescue – Morc3 wild type rescue, ATP.h – ATP hydrolysis mutant, ATP.b – ATP binding mutant, CW – CW mutant

